# G-Quadruplex-Protein Interactome at Human Gene Promoters

**DOI:** 10.1101/2025.01.01.630896

**Authors:** Rui-fang Duan, Jin-ping Zheng, Yu-hua Hao, Zheng Tan

## Abstract

DNA-protein interactions at gene promoters play a critical role in gene expression. The promoters of human cells are highly enriched in guanine-rich sequences, which can form four-stranded G-quadruplex (G4) structures. G4s are emerging as a distinct class of structure-based regulatory elements in gene regulation, and their interaction with proteins is essential for the role G4s play. Currently, our understanding of G4-protein interaction is mainly on a case-by-case basis, without systematic information. In this work, we examined the spatial occupancy of 1,183 human DNA-binding proteins, including transcription factors, histones and their modifying enzymes, around the consensus G4-forming region, G4(+), using data from the ENCODE project. We found that the G4(+), its immediate proximal side, and its distal side serve as three primary protein binding sites. Nearly all proteins are either enriched or depleted at these sites, likely due to competition, or in a spatiotemporal transition between the sites, resulting in different degrees of variation or persistence within or across cell/tissue types. Notably, histones were excluded from the proximal side of G4(+), and their binding to G4(+) was turned on and off by acetylation and methylation, respectively. Furthermore, the distal side is preferentially enriched for H3K23me2 and H3K4me2. Our experiments also revealed corresponding patterns of G4-protein interaction. Taken together, our results suggest a general role for G4s in dynamically defining and coordinating chromatin architecture and DNA-protein interactions at gene promoters for transcriptional regulation, a task that is unlikely to be accomplished by sequence-based DNA recognition.

Graphical abstract:A protein can be either enriched or suppressed at G-quadruplexes (G4s), at the proximal side of G4s, or enriched at the distal side of G4s in a 2 kb neighborhood. Regulatory roles of proteins may result from spatial transitions between the different binding regions.

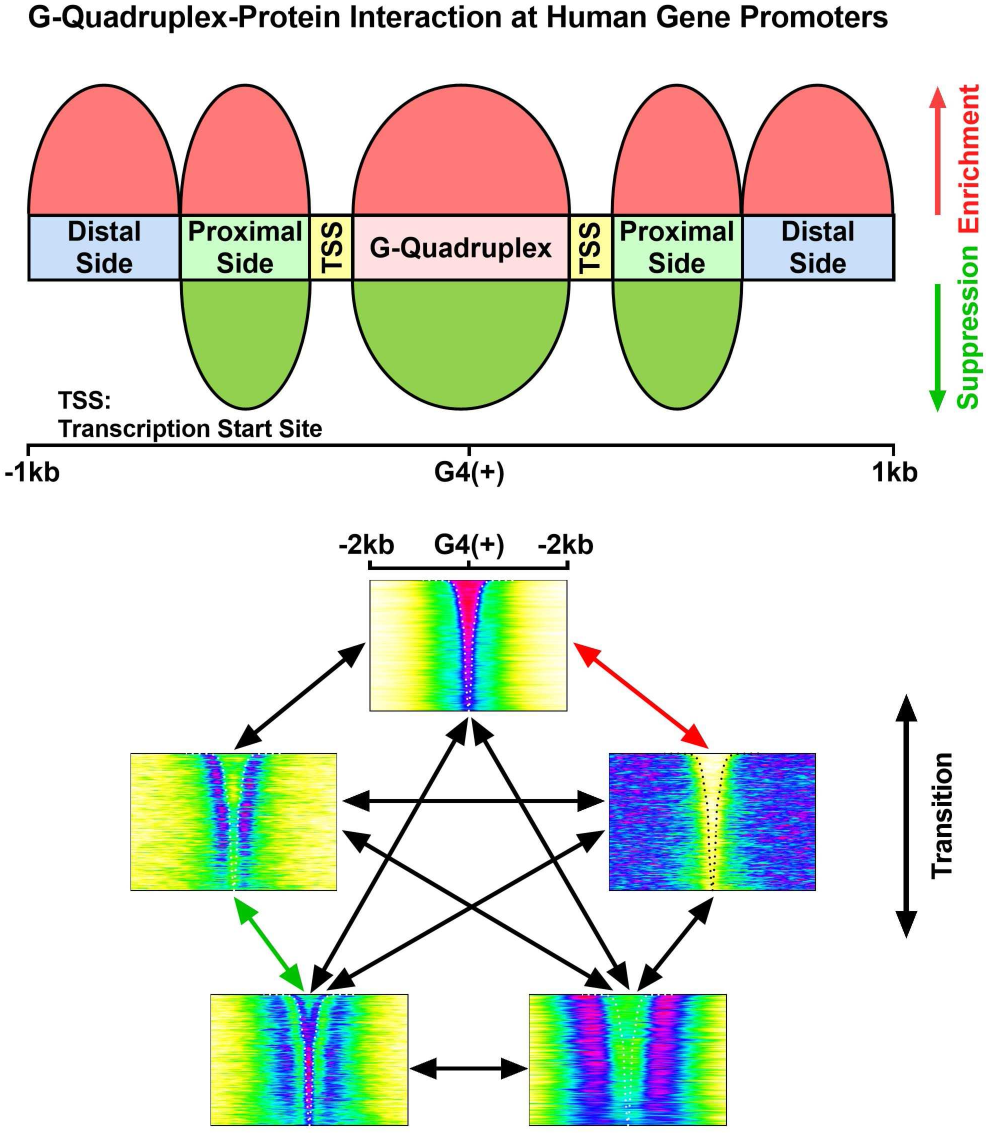

## Introduction

DNA encodes not only genes, but also gene regulation by sequence- or structure-based elements. DNA-protein interactions regulate many cellular processes through sequence-based recognition, such as the interaction of DNA with transcription factors (TFs). DNA also forms higher order structures. One such structure, the G-quadruplex (G4), is a four-stranded secondary structure that forms in guanine-rich nucleic acids (1). It has received increasing attention in recent years due to its important role in gene expression (2, 3). Putative G-quadruplex-forming sequences (PQSs) are abundant in the genomes of eukaryotic and prokaryotic cells. In particular, they are not randomly distributed, but highly enriched in the promoters of animal genes (4, 5), suggesting a general role in transcriptional regulation. Recently, G4s have been detected at PQS sites (6, 7), especially in promoters (6), in the human and animal genomes. According to a recently discovered mechanism of G4 formation in the eukaryotic genome (8), a single GG tract in a genome can form a G4 by recruiting G-tracts from RNA. As a result, a human genome can contain >150 million PQSs.

Human cells have >1500 TFs (9) and millions of transcription factor binding sites (TFBSs) (10). The interaction between TFs and TFBSs forms a complex and dynamic regulatory network in transcription. In addition to interactions via sequence-based recognition, G4-based regulation is expected to have a distinct and more diverse impact on the functionality of genomic DNA. Sequence-based DNA-protein interactions are more static on the DNA side, whereas G4-protein interactions are more dynamic, as a G4 unfolds and folds in response to cellular activities. In principle, a G4 can establish structure-based recognition, disrupt sequence-based interaction, physically impede protein translocation (11), segregate protein diffusion, inhibit DNA hybridization, and limit supercoiling wave transmission in DNA (12), to name a few.

Understanding the functionality of G4s in gene regulation requires information on how G4s interact with regulatory proteins, which has not been systematically studied except on a case-by-case basis with a limited number of proteins (13). To address this knowledge gap, we present a novel analysis of the interactome of G4s with such proteins in the human genome. We compiled the consensus of G4 formation from seven cultures of four human cell lines to examine the interaction between G4s and 1183 proteins, including transcription factors and chromatin remodelers, whose DNA binding data are available in the ENCODE project (10). By digitizing the protein occupancy around the G4 consensus, G4(+), in promoter regions, we identified distinct and well-defined DNA-protein interaction patterns across G4(+) that are preferentially adopted by proteins and are subject to spatiotemporal regulation. In particular, the interaction between histones and G4s is sensitively modulated by various modifications in a switch-like manner, allowing G4s to either recruit or repel a specifically modified histone protein. These features suggest that G4s play a general role in defining and coordinating chromatin architecture and diverse DNA-protein interactions to regulate gene expression.

## Results

### Consensus G4 formation

We aim to assess G4-protein interactions by examining the geometric presence of proteins on DNA around the consensus G4 site. Since DNA binding data of proteins have been collected on a large scale and are available from many sources, such as the ENCODE project (14), information on G4 formation at genomic loci is then critical to this goal. Recently, we identified G4 formation in four different living human cell lines using a small engineered G4 probe (G4P) protein that recognizes G4s with high specificity and affinity (6). These cell lines exhibited different landscapes of genomic G4 formation, and the number of G4 peaks detected differed significantly among them. Therefore, we hypothesize that the consensus G4 formation in these cell lines represents constitutive G4 structures that can be used to evaluate G4-protein interactions in these and other types of human cells.

We extracted the intervals of G4P peaks that were common to all four cell lines as consensus sites of G4 formation (Figure 1A). Because PQSs and G4 formation are both enriched around transcription start sites (TSSs) (6), we included only those peaks in the 2 kbp region upstream and downstream of the TSSs representing the promoters of human genes. The intervals were sorted by size in descending order (Figure 1B) and then concatenated to assemble a bed file (Figure 1C). By definition (Figure 1B), the bed file represents the regions of consensus G4 formation, denoted G4(+), surrounded by a region denoted G4(+/-), in which G4s may or may not form.

**Figure 1.**
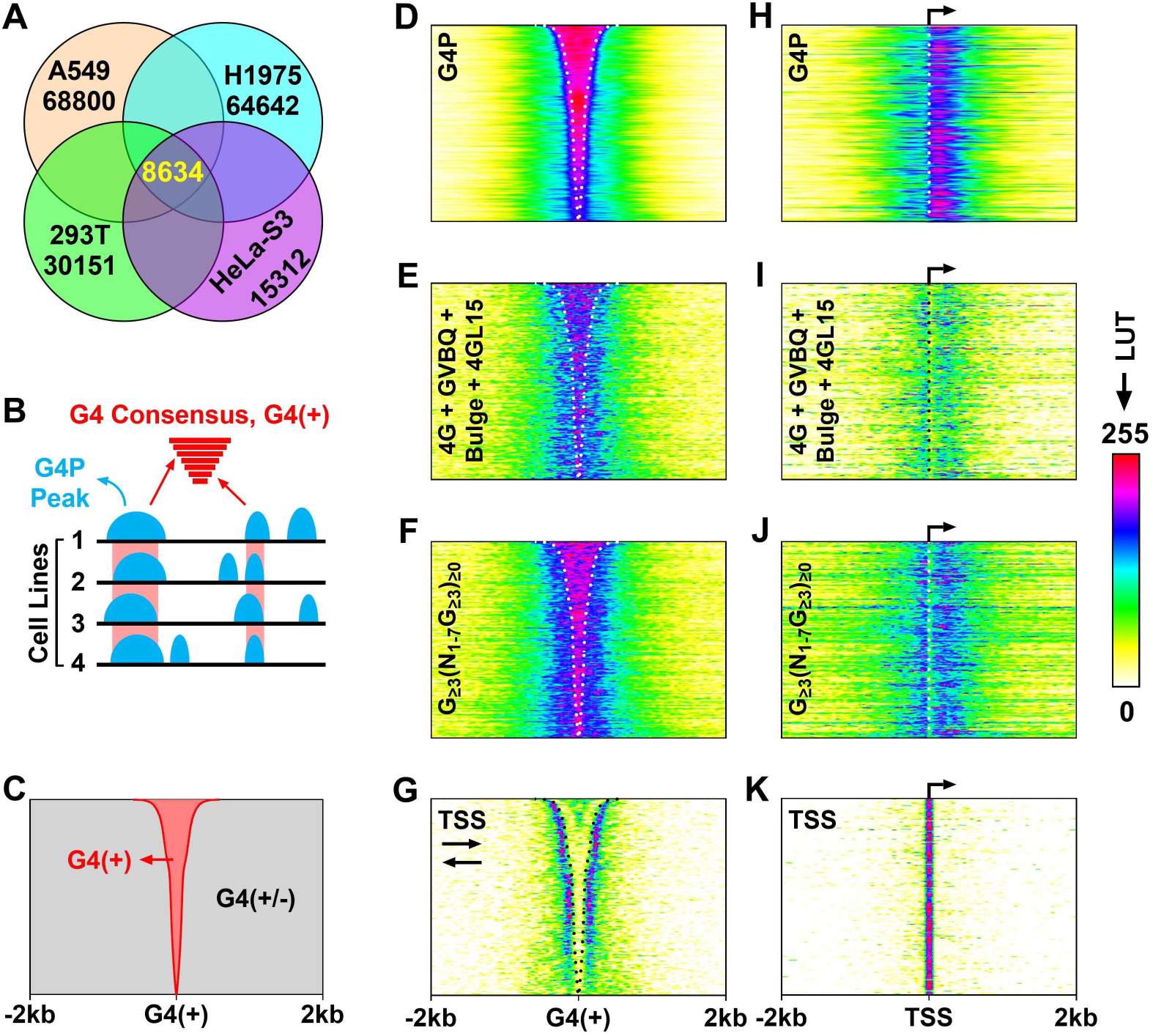
Consensus G4 formation and its association with putative G-quadruplex forming sequence (PQS) and transcription start site (TSS) in human gene promoters. (A) Overlap of G4P peaks in four human cell lines. (B) Extraction of G4 consensus, G4(+), from the common region of G4P peaks. (C) G4(+) regions sorted by interval size in descending order. (D) G4P occupancy across the G4(+) regions. (E-G) Distribution of canonical PQSs, contiguous G_≥3_ tracts, and TSSs across the G4(+) regions. (H) G4P occupancy across TSSs. (I-K) Distribution of canonical PQSs, contiguous G_≥3_ tracts, and TSSs across TSS. LUT, lookup table.

The G4(+) bed file allows analysis of the interaction of proteins or other genomic features with G4s by examining their occupancy around G4(+). Next, we calculated the occupancy of G4Ps and found that they were enriched in G4(+) as expected (Figure 1D). The same assessment was performed for PQSs with four consecutive G-tracts (6), resulting in a similar occupancy pattern (Figure 1E). Given our recent finding that motifs with as few as one G-tracts can form DNA:RNA hybrid G4s (hG4s) in the genome of eukaryotic cells (8), we also assessed the localization of such PQSs and found a similar but more significant enrichment in G4(+) (Figure 1F). This result not only further supports the formation of DNA:RNA hybrid G4s in eukaryotic cells (8), but also suggests a role for hG4s in DNA-protein interaction.

### Correlation between transcription and G4 formation

To determine the relationship between G4 formation and transcription, we calculated the localization of transcription start sites (TSSs) relative to G4(+). Interestingly, the TSSs showed a strong spatial association with G4(+) in its immediate proximal side (Figure 1G). This specific localization of TSSs to G4(+) suggests a direct involvement of G4s in transcription. To gain further insight, we identified TSSs that overlapped with G4(+) intervals by at least one nucleotide. The distribution of G4P, the two sets of PQSs, and the TSSs were then re-mapped to the identified TSSs. The results showed that the distribution of G4 formation, represented by G4P, was highly polarized and preferentially enriched on the downstream side of the TSSs (Figure 1H). The distribution of the two sets of PQSs also showed similar polarization (Figure 1, I-J) at the TSSs (Figure 1K). All these features suggest a deep involvement of G4s in transcription, for example, in defining transcription initiation.

### Prominent G4-protein interaction at G4(+)

To study G4-protein interaction, we downloaded 27448 bigwig files from the ENCODE database, each representing a protein from a sample of specific cell type or tissue, to calculate the protein occupancy of 1183 proteins around G4(+). The occupancy image of each sample was then compared to that of the reference G4P using the "compare" command of the Imagemagick software, which quantifies the difference between two images. The comparison returned a match score between 0 and 1 to describe the visual difference between the two images, where 0 means the two images are identical and 1 means they are completely different (Figure 2A, top versus bottom image). A cutoff of 0.7 identified proteins that showed similarity to the reference (Figure 2B). Using this criterion, more than half (>600) of the proteins were found to be enriched in G4(+), similar to G4P (Figure 2A, middle panel), suggesting a high affinity interaction between these proteins and G4s. The overall protein occupancy pattern qualitatively followed the density of PQSs (Figure 1, E-F), suggesting that such interactions are G4-specific.

**Figure 2.**
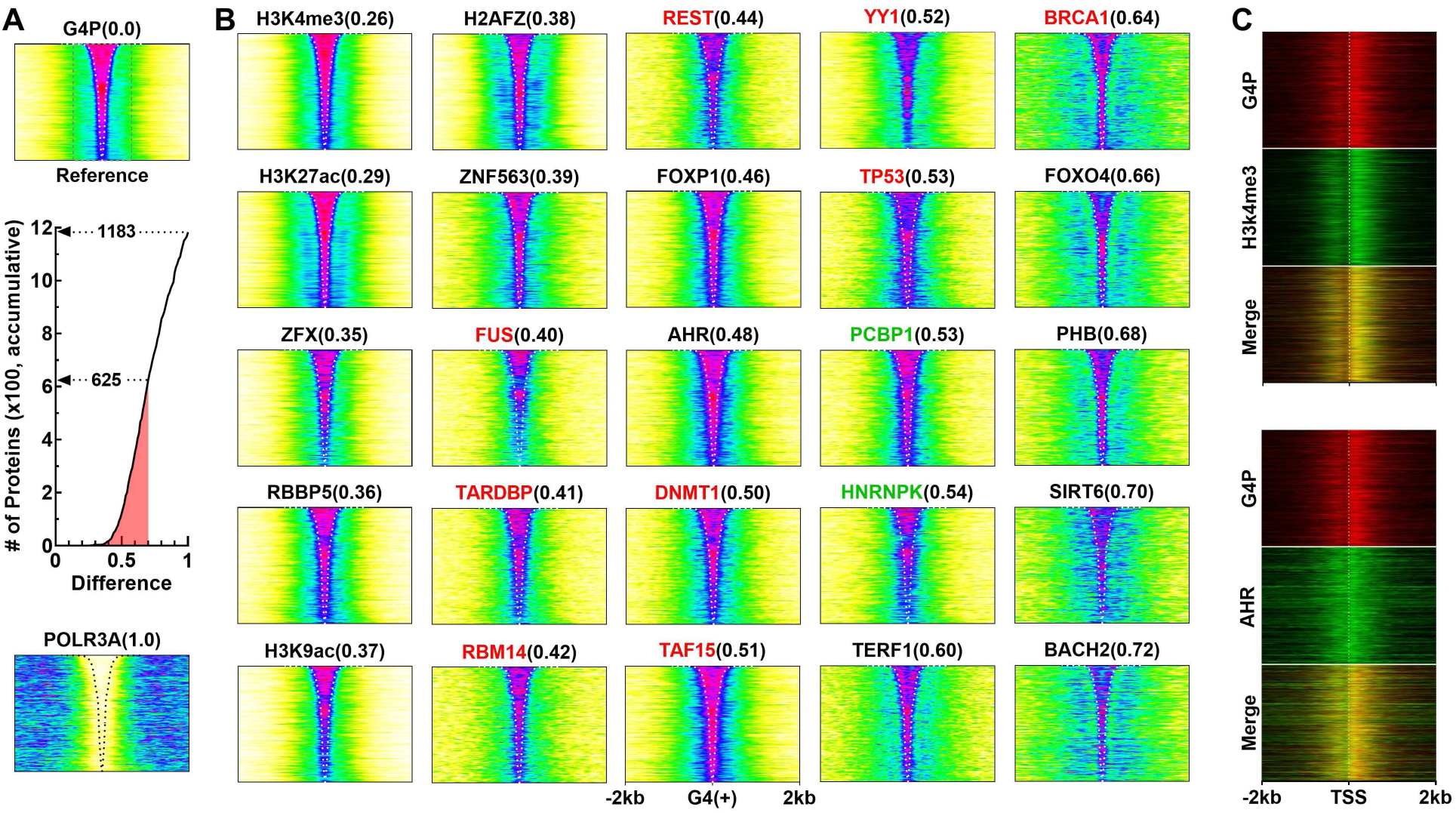
Protein occupancy across G4(+) compared to G4P enriched at the G4(+). (A) Number of proteins with a difference value from G4P in occupancy image less than or equal to the indicated value. Top image is reference in which the dashed box indicates the core region used for image comparison. (B) Representative occupancy of proteins across G4(+). Names in red are known G4-binding proteins and those in green are C-rich DNA binding proteins. Numbers in parentheses indicate visual differences from the reference. Dashed lines indicate boundaries of G4(+). (C) Occupancy of G4P, H3K4me3, and AHR across transcription start site (TSS).

Among the identified proteins, some are known G4-binding proteins (15, 16) and a few are known to bind C-rich sequences (17, 18) (Figure 2B, names in red and green, respectively). In particular, H3K4me3 from the Peyer’s patch tissue showed an occupancy virtually identical to that of G4P (Figure 2B), suggesting it being a strong G4 binder. To investigate the correlation between H3K4me3 and G4 in transcription, we re-mapped H3K4me3 and G4P around the TSSs. The results showed that both H3K4me3 and G4 formation were enriched on both sides of the TSS, with a greater magnitude on the downstream side, as indicated by the yellow signal in the merged image (Figure 2C, three panels at top). As a chromatin modifier associated with the activation of gene expression, H3K4me3 is found in the promoter regions of active genes (19). Therefore, the observed enrichment of H3K4me3 at G4(+) clearly indicates an involvement of G4 in transcriptional regulation. A similar result was obtained with aryl hydrocarbon receptor (AHR) (Figure 2C, lower panels), a transcription factor with roles in detoxification, development, immune response, chronic kidney disease and other syndromes (20). These results strongly suggest that G4s may play a role in recruiting proteins to TSSs for transcription.

Given the large number of DNA-binding proteins, competition among them for common target sites is expected. Accordingly, some proteins were depleted at G4(+) (Figure 3). CEBPZ, a transcriptional activator involved in cellular responses to environmental stimuli (21), was found so in the HepG2 cell line. Using it as a reference (Figure 3A, top image), the presence of 186 proteins was found to be suppressed to varying degrees at G4(+) (Figure 3B). In principle, these proteins could be excluded from G4(+) by stronger G4 binding activities, or they may not recognize G4 and therefore occupy only the G4-free regions. The mutual exclusion between the distribution of G4P and that of CEBPZ and DDX20 across the TSS further demonstrated that the latter two proteins were avoided in the presence of G4 (Figure 3C).

**Figure 3.**
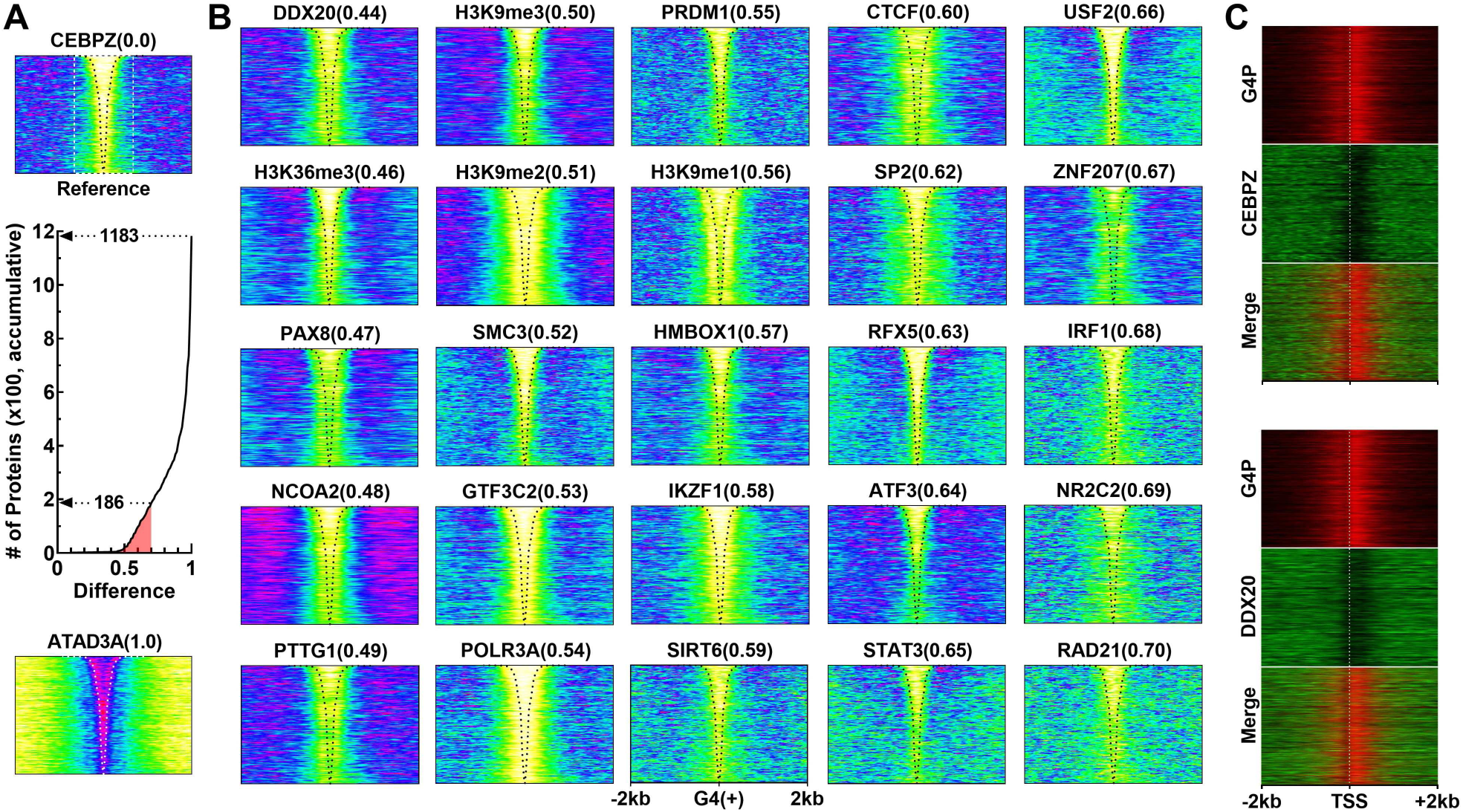
Protein occupancy across G4(+) compared to CEBPZ depleted in G4(+). (A) Number of proteins with a difference value from CEBPZ in occupancy image less than or equal to the indicated value. (B) Representative occupancy of proteins across G4(+). (C) Occupancy of G4P, CEBPZ, and DDX20 across TSS. Other annotations are the same as in Figure 2.

### Prominent G4-protein interaction next to G4(+)

In addition to the enrichment at G4(+) (Figure 2), a significant enrichment in a well-defined region adjacent to G4(+) was observed for over 270 proteins (Figure 4A), using SP1 from the HepG2 cell line as a reference. SP1 is a well-characterized transcription factor known for its ability to bind to GC-rich DNA sequences and its role in a variety of essential biological processes (22). Its enrichment on the proximal side of G4(+) was G4-dependent, as its intensity decreased with increasing distance from G4(+) (Figure 4B), similar to the behavior of G4 formation and PQSs (Figure 1, D-F). Interestingly, this region (Figure 4B) was tightly co-localized with the TSS (Figure 1G), suggesting that such enrichment may be involved in transcription initiation. Remapping of SP1 and another protein, DRAP1, over the TSS further revealed that these two proteins were predominantly enriched on the upstream side of the TSS, likely on the G4-free regions, as this area showed significantly less signals from G4P (Figure 4C).

**Figure 4.**
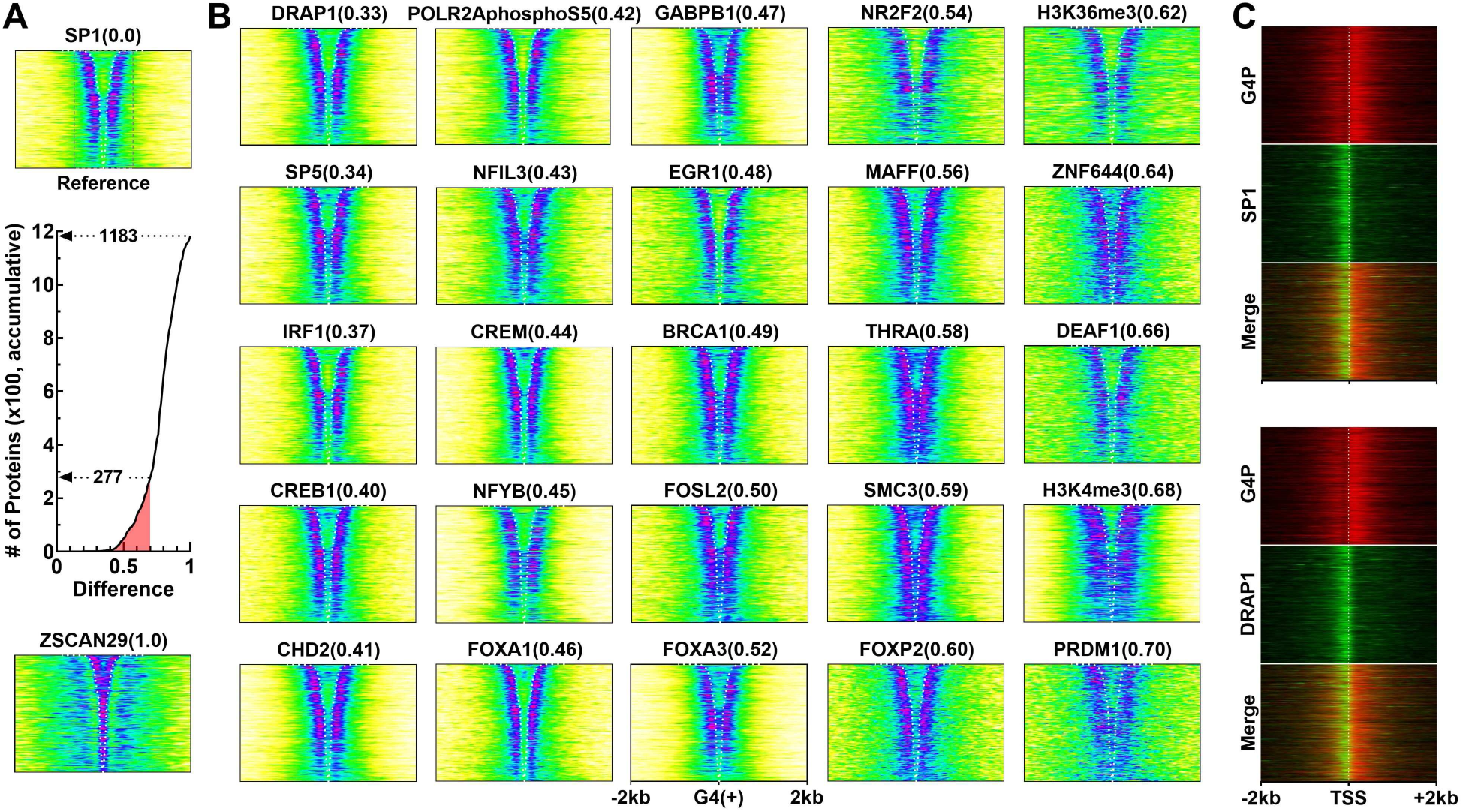
Protein occupancy across G4(+) compared to SP1 enriched at the G4(+) proximal side and near TSS. (A) Number of proteins with a difference value from SP1 in occupancy image less than or equal to the indicated value. (B) Representative occupancy of proteins across G4(+). (C) Occupancy of G4P, SP1, and DRAP1 across TSS. Other annotations are the same as in Figure 2.

In contrast to the enrichment, a suppression on the G4(+) proximal side was found for H2AFZ (Figure 5A) from the H1 cell line, a variant of the histone H2A that plays a role in chromatin regulation by affecting nucleosome structure and function to influence gene expression, DNA repair, replication, and chromosome stability (23). Similar suppression was found for over 400 other proteins (Figure 5B), using the H2AFZ as a reference. The overall distribution in the remaining regions largely followed the gradient of G4 formation and PQS (Figure 1, D-F), suggesting that they were associated with G4s, but were excluded on the G4(+) proximal side by stronger competition from other G4 binders. Furthermore, the suppression on the proximal side of G4(+) was often accompanied by suppression in the inner region of the larger G4(+) sites, resulting in a "Y-shaped" protein enrichment of varying degrees in G4(+). In contrast to the results shown in Figure 4C, these proteins were suppressed immediately upstream of the TSSs, suggesting that they (Figure 5C) may have a role opposite to that of the proteins in Figure 4.

**Figure 5.**
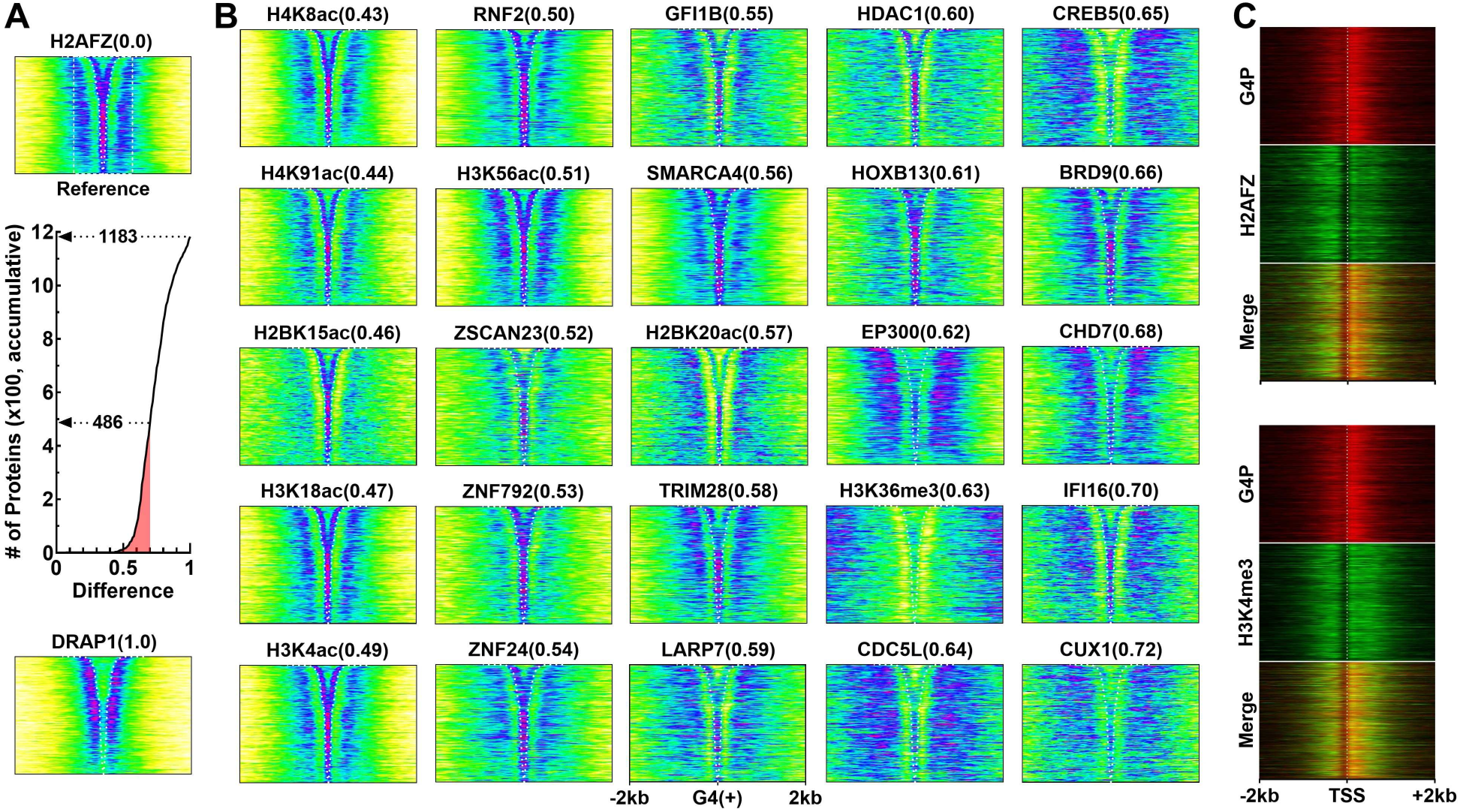
Protein occupancy across G4(+) compared to H2AFZ depleted at the G4(+) distal side G4(+) and near TSS. (A) Number of proteins with a difference value from H2AFZ in occupancy image less than or equal to the indicated value. (B) Representative occupancy of proteins across G4(+). (C) Occupancy of G4P, H2AFZ, and H3K4me3 across TSS. Other annotations are the same as those in Figure 2.

### G4-protein interaction distal to G4(+)

In addition to the enrichment at G4(+) and its proximal side, a broader enrichment was also observed at the distal side of G4(+) for more than 160 proteins (Figure 6), using H3K4me2 from the A549 cell line as a reference (Figure 6A, top image). Like the H3K4me3, H3K4me2 is also an epigenetic mark that regulates gene transcription. In this case, a much weaker signal was detected at G4(+) and its proximal surroundings (Figure 6A), likely due to competition from other G4-binding activities in these two regions. Many proteins in this group (Figure 6B) were distributed similarly to those shown in Figure 5B. Remapping of the representative H3K4me2 and H3K27ac across the TSSs revealed that they were enriched at where G4 formation was weak (Figure 6C), suggesting that their interaction with DNA may be modulated by G4.

**Figure 6.**
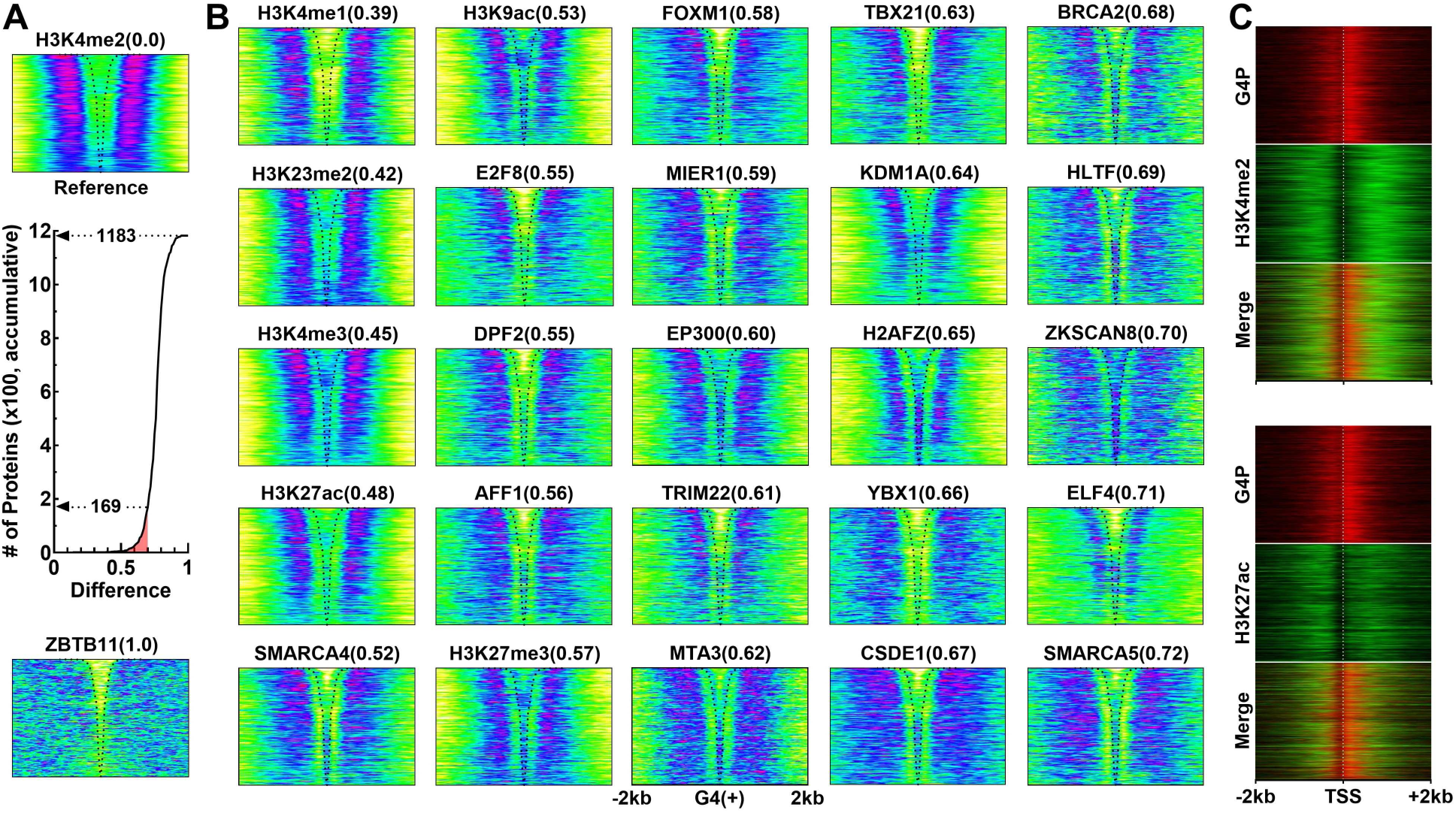
Protein occupancy across G4(+) compared to H3K4me2 from A549, which is enriched at the G4(+) distal side. (A) Number of proteins with a difference value from H3K4me2 in occupancy image less than or equal to the indicated value. (B) Representative occupancy of proteins across G4(+). (C) Occupancy of G4P, H3K4me2, and H3K27ac across TSS. Other annotations are the same as in Figure 2.

### Cell-type specific protein-G4 interaction

Cell type-specific gene expression is the most important aspect of development, differentiation, and pathology. We examined protein occupancy in a cell type-specific manner and found that proteins exhibited varying degrees of consistency or variation within or across cell types. Extreme persistence was observed for NRF1, which showed nearly identical proximal G4(+) side enrichment in all samples from the seven cell types (Figure 7A, Figure S1). This consistency across cell types was also observed for PHF8, which showed G4(+) enrichment in all samples from A549, H1, HepG2, and K562 cells (Figure S2), for HBTF in four, and for CHD2 (Figure S3 and Figure S4) in seven cell types.

**Figure 7.**
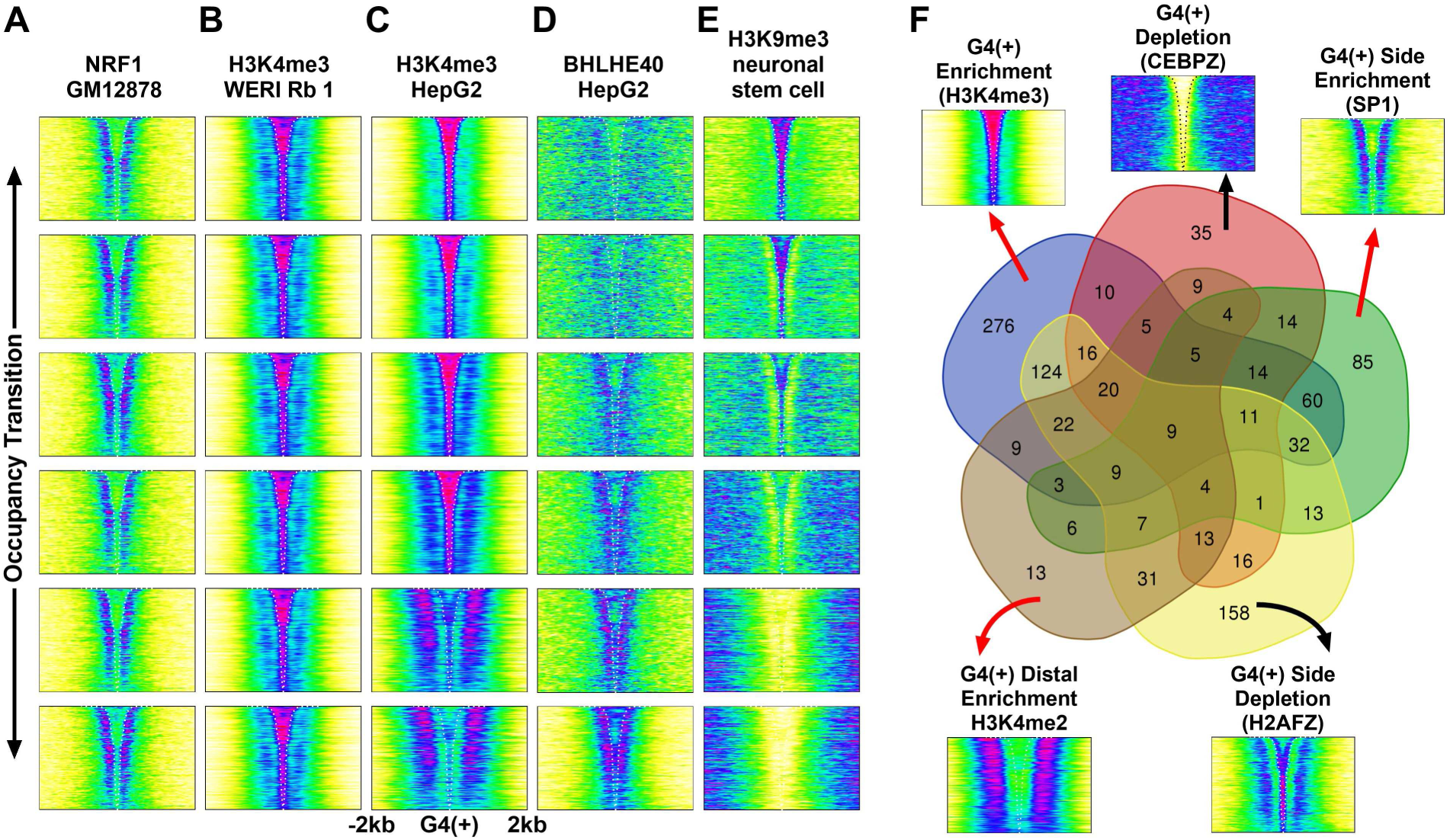
Consistency and variation of protein occupancy across G4(+). (A-E) Representative occupancies of proteins in samples from the same cell type. (F) Venn diagrams of representative occupancy patterns observed in 1053 out of 1183 proteins.

Consistency was more often observed between samples of the same cell type. For example, H3K4me3 showed dominant enrichment at G4(+) in all six samples of WERI Rb 1 cells (Figure 7B). H3K27ac showed dominant enrichment at G4(+) in all 83 samples of A549 cells (Figure S5), and H3K4me1 was depleted at G4(+) and its vicinity in all 60 samples of A549 cells (Figure S6). In these cases, inter-cell type variations were also frequently observed. For example, H3K4me3 showed a predominant enrichment at G4(+) in all lower leg skin, BJ, and coronary artery samples, but a suppression at the G4(+) proximal side accompanied by an enrichment at G4(+) and its distal side in all BE2C, AG10803, and kidney cell samples (Figure S7). Additional examples of consistency within and variation among cell types are shown in Figures S8-S18.

On the other hand, some proteins showed drastic changes in occupancy within the same cell type. For example, H3K4me3 in HepG2 cells displayed a transition from dominant enrichment at G4(+) to dominant enrichment at the G(+) distal side (Figure 7C). For BHLHE40 in HepG2 cells, a gradual development of enrichment towards the G4(+) proximal side was observed from a more diffuse distribution (Figure 7D). The enrichment of H3K9me3 from the neural stem cells at G4(+) was gradually reversed, ultimately resulting in synergistic repression of H3K9me3 at G4(+) and its surrounding region (Figure 7E). These results demonstrate a dynamic spatiotemporal modulation of G4-protein interaction for gene regulation. Additional examples of variation in different cell types are shown in Figure S19.

Figure 7F provides a general assessment of the consistencies and variations in protein spatial occupancy, covering 1053 (89%) of the 1183 proteins and 16012 (58%) of the 27448 samples examined. Since each source dataset was obtained from a large population of cells, well-defined patterns of protein distribution indicated that they were statistically significant features. As we can see, a protein can have a distribution that belongs to more than one category (e.g., Figure 7, C-E). The proteins or samples that were not covered did not show typical features on the overall scale. They often showed a diffuse or mixed and occasionally random distribution pattern, which may represent transitional intermediates between typical patterns or a mixture of typical features in a heterogeneous cell population.

### Interplay among proteins at and near G4s

Our results revealed two main types of G4-protein interactions, represented by the enrichment and suppression of proteins in G4(+) (Figures 2, 3) and on its proximal side (Figures 4, 5). The suppression or depletion of proteins can be explained by competition between proteins at these sites. In Figure 8, we provide four hypothetical examples to illustrate how a stronger competitor may exclude a protein from the two regions.

**Figure 8.**
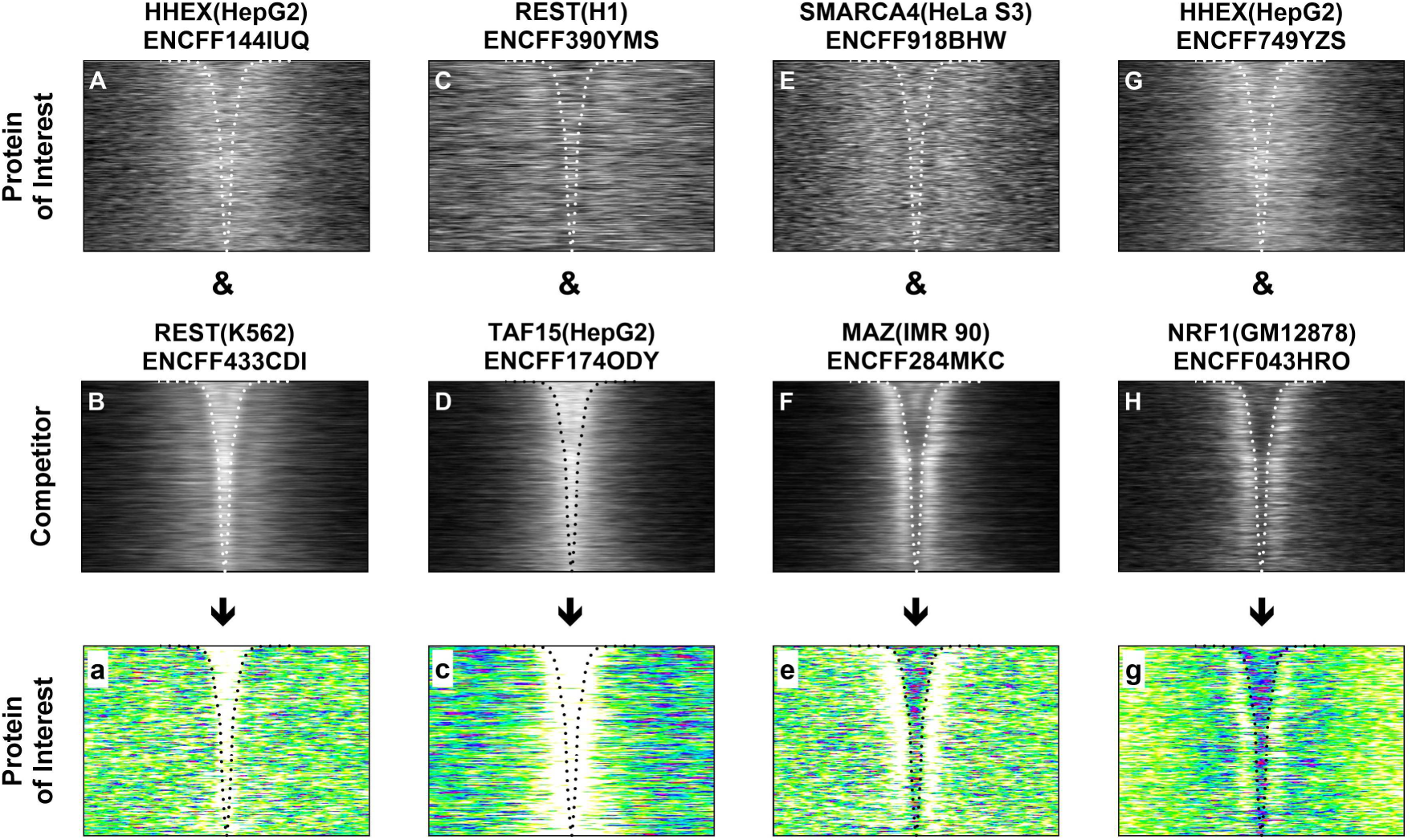
Suppression/depletion of proteins in G4(+) and on the G4(+) proximal side as a result of a hypothetical competition between two proteins. The image of a competitor (second row) was superimposed on that of the protein of interest (first row) using ImageJ (https://imagej.net/ij/index.html) in the "Subtract" transfer mode, and the resulting image was then converted to a color image (third row) using the custom lookup table shown in Figure 1. Cell lines are shown in parentheses with the ID below.

### Spatial preference of protein occupancy

Our results also showed that some proteins prefer to adopt a specific occupancy pattern around G4(+). Proteins in the ENCODE database can be divided into two groups: 1) transcription factors and 2) histones and their modifying enzymes. To assess their spatial preferences, we examined the percentage of samples for each protein that adopted a particular distribution pattern represented by the reference proteins (Figure 2-6, panel A). The results for the transcription factors in Figure 9 identified a few strong G4-interacting proteins that were enriched in G4(+), including BCL3, CHD1, JUN, NONO, SMC3, and ZFX (Figure 9A). In comparison, more proteins such as CHD2, CREB1, E2F4, ELF1, FOS, GABPA, MAX, MAZ, NRF1, RFX5, RXRA, SP1, SRF, TBP, USF1, and ZNF143 were preferentially enriched at the G4(+) proximal side near the TSS (Figure 9C). Depletion at G4(+) and its proximal side and enrichment at G4(+) distal side were much less frequent (Figure 9, B, D, and E).

**Figure 9.**
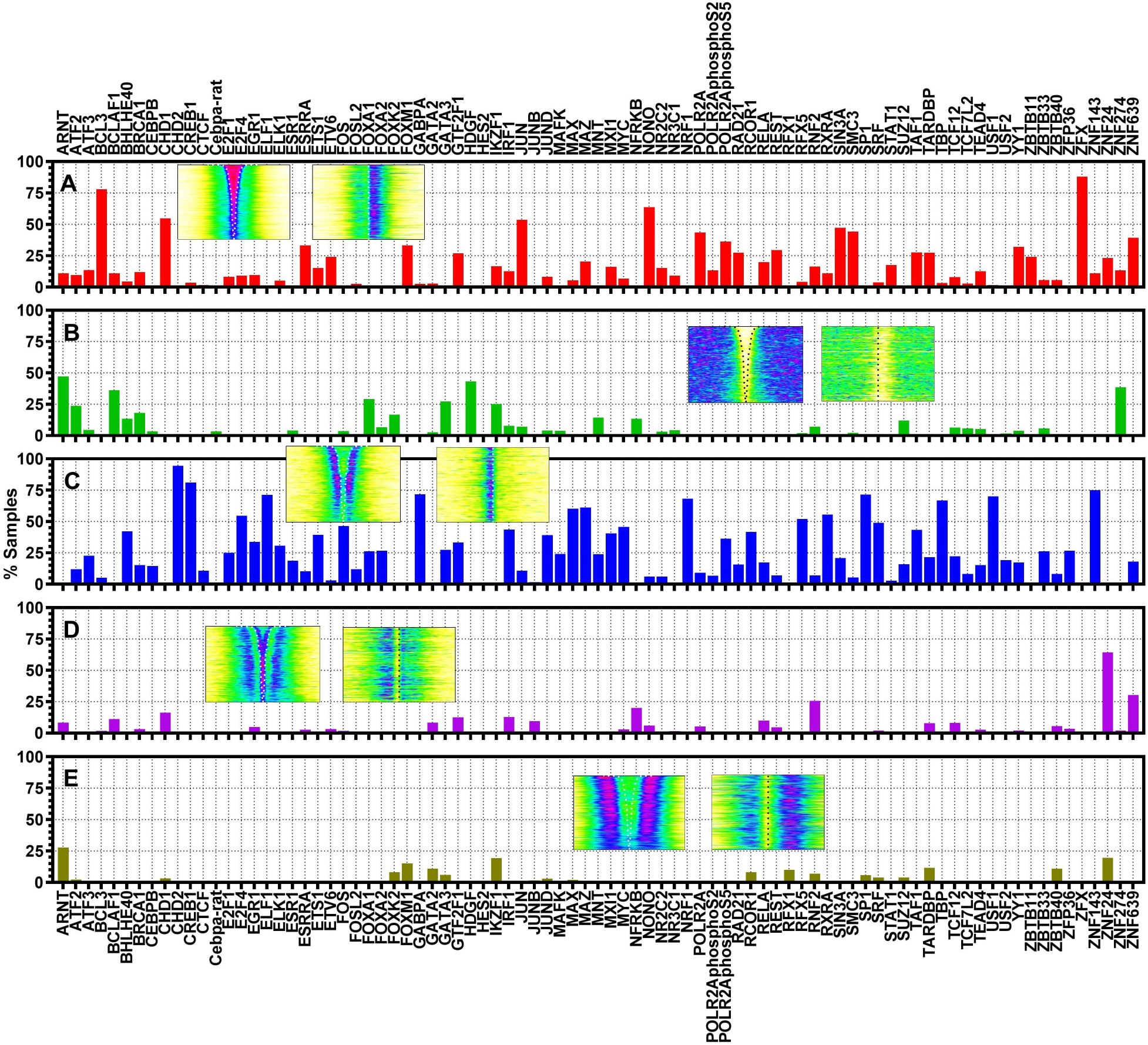
Spatial preference of 85 proteins, expressed as percentage of samples with a visual difference of ≤0.7 to the reference in Figure 2-6. The occupancy images of the references are inserted in the order of H3K4me3, CEBPZ, SP1, H2AFZ, and H3K4me2 (from top to bottom) around G4(+) and TSS (from left to right, dotted lines), respectively. Proteins with fewer than 30 samples are ignored.

Nearly half of the samples were for histones with various modifications that are responsible for packaging chromosomal DNA into nucleosomes (24). These epigenetic features modulate chromatin organization and gene expression in which G4s play a role (25). The results for histones and their modifying enzymes are shown in Figure 10, with several unique features. First, virtually all acetylated histones showed a strong preference for enrichment at G4(+) (Figure 10A), mostly accompanied by depletion at the G4(+) proximal side (Figure 10D), indicating their strong binding to G4s. Second, methylated histones were preferentially depleted at G4(+) (Figure 10B), suggesting that methylation deprives histones of their ability to bind G4s. Third, the effect of methylation on histone H3 lysine 4 (H3K4) is an interesting exception, with a transition from depletion to enrichment at the G4(+) as the methylation increased from 1 to 3 (Figure 10, A and B, asterisk). Consider BLaER1 cells as an example. While most of the mono-methylated H3K4 was completely depleted at G4(+) in these cells in contrast to the surrounding regions (Figure S16), the di-methylated H3K4 started to enrich at G4(+) along with a significant concentration at the G4(+) distal side (Figure S17). The tri-methylated H3K4 then became predominantly enriched at G4(+), with decreasing occupancy as it moved away from G4(+) (Figure S18). Fourth, all histones avoided enrichment exclusively on the G4(+) proximal side (Figure 10C). Fifth, H3K23me2 and H3K4me2 were exceptional in that they both tended to be highly enriched on the G4(+) distal side (Figure 10E).

**Figure 10.**
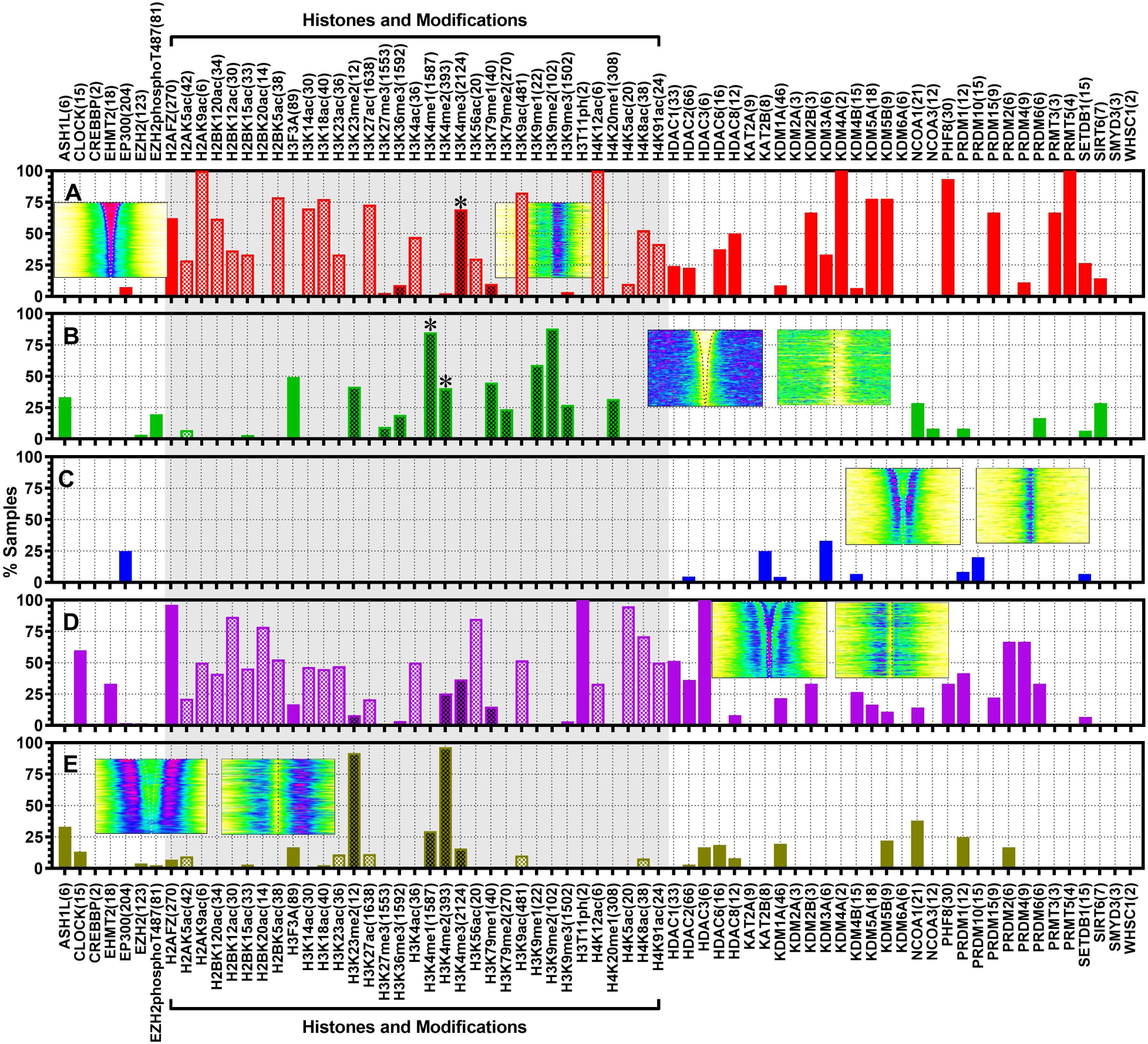
Spatial preference of histones and histone modifying enzymes, expressed as percentage of samples with a visual difference of ≤0.7 to the reference in Figures 2-6. The number in parentheses indicates the number of samples for each protein to alert potential statistical variation for proteins with small sample sizes. White-dotted bar, acetylation; black-dotted bar, methylation. Image inserts are the same as in Figure 9.

### Recognition element in G4-Protein interaction

The formation of G4 at genomic loci results in several structural features. In addition to G4, the cytosine-rich (C-rich) complementary strand can fold into an i-motif or remain relaxed at acidic intracellular pH (26). These structures, including G4, i-motifs, relaxed C-rich strands, and their interfaces with flanking regions, may serve as potential targets for protein recognition (Figure 11A). To elucidate such molecular events, we performed competitive electrophoretic mobility shift assays (EMSAs) on three representative proteins, EGR1, PCBP1, and NONO, with an appropriate DNA substrate.

**Figure 11.**
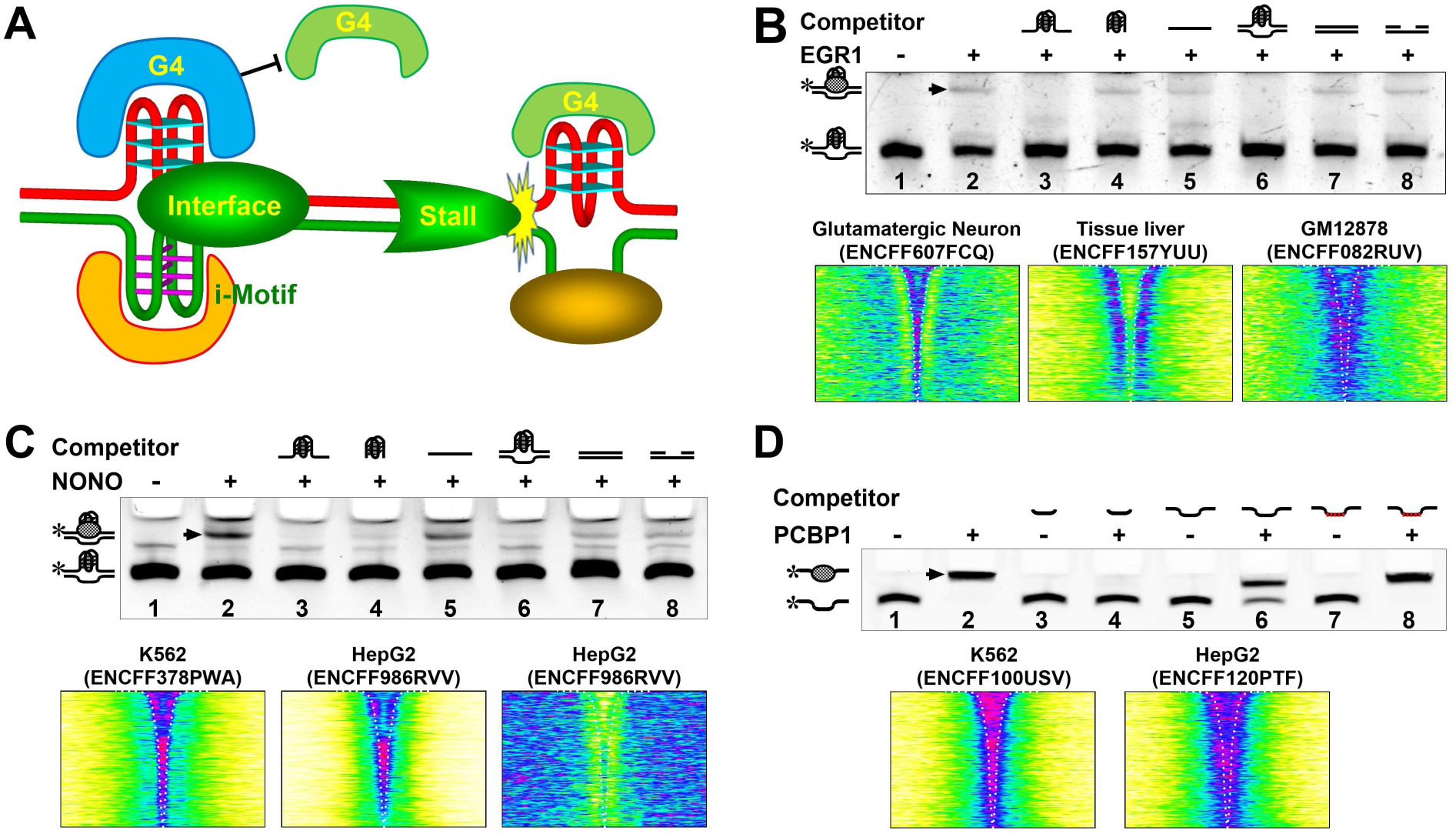
Expected effects of G4 formation on proteins and examples of DNA-protein interactions examined *in vitro*. (A) Illustration of potential DNA-protein interactions at G4 sites. (B-D) Binding of EGR1, NONO, and PCBP1 to G4s revealed by EMSA and their spatial occupancy across G4(+). Arrowhead indicates DNA-protein complex.

As shown in Figure 11B, EGR1 bound to a dsDNA containing a G4 in the middle (lane 2). This binding required both the G4 and flanking sequences, as evidenced by the disappearance of the shift band when the binding reaction was performed in the presence of an unlabeled single-stranded (lane 3) or double-stranded (lane 6) G4 DNA competitor with flanking sequences. In contrast, a naked G4 (lane 4) or other non-G4 DNAs did not show such an competition (lanes 5, 7-8). Representative distributions of EGR1 showed that the protein was enriched within G4(+) in glutamatergic neurons, resulting in a "Y-shaped" pattern, suggesting that the flanking sequence was also involved in G4 recognition, so that the protein avoided the inner region of G4(+). This result was consistent with the EMSA (lane 4), where a naked G4 did not compete with the flanked G4. In contrast, EGR1 in the liver sample switched to the outside of G(+) next to the G4(+) boundary, suggesting that recognition to the flank was maintained. In the GM1287 cell line, EGR1 appeared to be in a transition between the two previous patterns, with a more diffuse concentration at the G4/flank interface. In all these cases, interface recognition may play a role.

Another G4-binding protein, NONO, showed more efficient binding to the G4 DNA (Figure 11C, lane 2). Binding was prevented by a single-stranded (lane 3) or double-stranded (lane 6) DNA with a G4 in the middle. A naked G4 did not completely prevent the binding (lane 4), suggesting that binding of NONO to G4 was facilitated to some extent by flanking strands. A random single-stranded DNA slightly reduced the binding (lane 5), and a greater reduction was seen with the other non-G4 competitor (lanes 7 and 8). These features suggest that the recognition of NONO for G4s was not as specific as that of EGR1. For both EGR1 and NONO, the weak shift band implies low affinity binding to TFBSs, a common and functional aspect of transcription factor activity (27).

We also tested a C-rich DNA binding protein, poly(C)-binding protein 1 (PCBP1), which binds to the 5’-UUCCCUCCCUA-3’ RNA sequence (28). This protein bound strongly to single-stranded C-rich DNA with flanks (Figure 11D, lane 2). The binding was prevented by an unlabeled C-core sequence, demonstrating that the flanks were not necessary for recognition (lane 4). Adding flanks to the C-core reduced the binding of the labeled DNA (lane 6) by ∼1/3, possibly due to a slow dissociation of the unlabeled DNA from the DNA/PCBP1 complex. In contrast, random DNA showed no effect (lane 8). The spatial distribution of PCBP1 was available from two cell lines, which showed a dominant (K562) and a more diffuse (HepG2) enrichment in G4(+), respectively, consistent with the ability of PCBP1 to bind the C-rich strands opposite to G4s.

## Discussion

As a comprehensive G4-protein interactomic study, our results revealed three distinct major regions in human gene promoters: G4(+), G4(+) proximal side, and G4(+) distal side (Figure 7F, red arrowheads), in close proximity to TSSs (Figures 2, 4, 6, column C), where proteins bind DNA in well-defined spatial patterns relative to G4s. Because these three regions are shared by almost all proteins, competition between proteins occurs in the binding regions, leading to suppression of protein occupancy (Figures 3, 5). Thus, complex interactions between proteins occur in a spatiotemporally regulated manner across cell/tissue types (Figure 7), resulting in the transition of proteins between regions, presumably as a means of functional regulation. As our study included transcription factors, histones, and their modifying enzymes, these features reflect a critical role of G4s in orchestrating chromatin architecture and coordinating overall DNA-protein interactions in transcriptional regulation.

On the DNA side, G4(+) represents a polarized composition of G4s across TSSs (Figure 1D versus Figure 1H), with a greater amplitude on the downstream side than on the upstream side. G4s can bind more than half of the proteins (Figure 2A), either directly or indirectly via intermediates. Furthermore, the G4(+) proximal side (Figure 4A, top panel), which represents the immediately upstream side of TSSs with low probability of G4 formation (Figure 4C), served as another set of binding sites for nearly 300 proteins. In particular, a region distal to G4(+) and TSS was specifically recognized by both H3k23me2 and H3k4me2 (Figure 6A, top panel). On the protein side, competition between proteins (Figure 8) at the first two sets of binding sites satisfactorily explains the suppression of proteins at these sites (Figures 3, 5), suggesting a complex interplay among proteins near or at the G4(+). In summary, proteins intend to adopt one of the five distinct spatial states in association with G4s or in transition between them.

Of all the non-histone proteins analyzed, more proteins attempted to bind to the G4(+) proximal side than to the G4(+) (Figure 9, panel C versus A). Suppression of proteins at these two sites occurred much less frequently (Figure 9, panels B and D). In contrast, histone proteins generally behaved in the opposite manner, with two unique features. First, they mostly recognized G4(+) (Figure 10A), but avoided the G4(+) proximal side (Figure 10C). Second, the binding of histones to G4s was determined by the acetylation/methylation modification of lysines. In general, acetylated histones bind G4s (Figure 10A), whereas methylated histones are prevented from binding (Figure 10B). Interestingly, H3K4 showed a switch-wise binding behavior depending on the level of methylation (Figure 10, A versus B). H3K4me1 was strongly excluded from G4(+), and this exclusion was reduced for H3K4me2 after the addition of an extra methyl group. H3K4me3 then became a strong binder of G4. Histone modifying enzymes also showed a spatial preference. Most of them could bind to G4(+) (Figure 10A), and a few could bind to the G4(+) proximal side (Figure 10C).

Transcription is closely linked to the accessibility of chromatin, the complex of DNA and histone proteins within the nucleus of eukaryotic cells (29), where binding of TFs to TFBSs plays a more direct role in transcriptional regulation (9). TFBSs and PQSs are both enriched at promoters in the human genome, particularly in the vicinity of TSSs (30), providing a geographic basis for G4s to interact with TFs. In contrast to sequence-based recognition, which is largely static, G4s offer a more dynamic and versatile regulation due to their structural diversity (31), physical stability (32), dynamic nature (33), and ability to respond to the environment (5), which are sensitive to multiple intrinsic and environmental factors (34). The recently identified formation of DNA:RNA hybrid G4s in the eukaryotic genome (8) introduces further structural diversity resulting from the different ratios of DNA:RNA G-tracts in G4 formation and the ability to respond to RNA levels due to the involvement of RNA.

Since the three recognition sites are shared by almost all proteins examined, we infer that the role a protein plays in transcription is defined by its spatial distribution controlled by G4 formation. While more detailed studies are needed to unravel the underlying mechanism, several events may be involved in such processes. A G4 can be directly bound by proteins with varying degrees of affinity. In addition, it is well documented that G4s act as roadblocks to protein translocation (11, 35), which would be expected to affect the movement of any protein on DNA. The landing of a protein on DNA is random, followed by stochastic sliding along the DNA until it finds its target (36). A G4 can slow down, pause, or stop protein translocation, depending on its physical stability and the G4-resolving activity in cells. In addition, two or more G4s tethered to each other in a DNA strand can also trap proteins between them. In this way, G4s may act like speed bumps or traffic lights to control the localization and distribution of proteins and coordinate their genomic activities, organizing a complex interplay of proteins with DNA in a spatiotemporally regulated manner.

The association of robust G4 formation with TSSs (Figure 1H) and the ability of G4s to modulate histone and TF binding (Figures 9, 10) in promoters support a general role for G4s in defining and coordinating the interaction of DNA with transcription factors and histones, providing a molecular basis for a role of G4s in orchestrating chromatin architecture and mediating DNA-protein and protein-protein interactions in transcriptional regulation. Technically, the G4(+) was assembled to facilitate the analysis of G4-protein interactions with better spatial resolution than alignment at TSSs. However, the observation made in this approach may also occur for other short-lived G4 sites whose interaction with proteins occurs in a much shorter time window that is difficult to track.

## Materials and Methods

### Genome sequence and gene features

These files were downloaded from the UCSC genome browser website (genome.ucsc.edu).

### Consensus of G4 formation

G4P narrow peaks were identified using the Macs2 software (37), version 2.2.6, with parameters set to -qvalue 0.001, -keep-dup 1, and default values for the other parameters. For the A549, 293T, and NCI-H1975 cell lines, reproducible G4P peaks were derived from two biological replicates using irreproducible discovery rate (IDR) analysis with default parameter settings (38). The resulting peak files and that for HeLa-S3 were then used to identify the common intervals of G4P peaks shared by all four cell lines using the Bedtools Multiinter software (39), version 2.29.2. After eliminating those lying outside the ±2kb region of transcription start sites (TSSs), 8634 G4P peak intervals were obtained and compiled into a bed file. The bed file was sorted by peak interval size in descending order to represent the consensus of G4 formation, referred to as G4(+).

### Identification of PQSs in genome and their distribution around genomic regions

Three sets of PQS motifs were identified from the genome sequences. The PQSs with four G-tracts, i.e., 4G, 4GL15, Bulge, and GVBQ, were identified as described (6). The PQSs with one and up to three G_≥3_ tracts were identified using a regular expression of G{3,}(.{1,7}?G{3,}){0,2} with a standalone program as described (4). The PQSs capable of forming DNA:RNA hybrid G4s or canonical G4s were identified using a regular expression of G{3,}(.{1,7}?G{3,}){0, } with a standalone program as described (4). The resulting PQS bed files were converted to bedgraph files using the Bedtools genomecov software (39), version 2.29.2, and then converted to bigwig files using the bedGraphToBigWig software from UCSC (http://genome.ucsc.edu/). Their distribution at genomic regions was calculated as described (6).

### Score data of protein occupancy from ENCODE

Bigwig files representing fold changes in protein occupancy signal were downloaded from the ENCODE website (www.encodeproject.org). As of October 5, 2023, a total of 27448 bigwig files, in output type of fold change over control, were obtained for 1183 proteins. Their distribution at genomic regions was calculated as described (6).

### G4P ChIP-Seq data analysis

Clean paired-end G4P-ChIP sequencing data in fastq format for the cell lines A549, 293T, NCI-H1975, and HeLa-S3 were obtained from our previous study (6), which can be downloaded from the GEO database at NCBI under accession number GSE133379. DNA reads were mapped to the human hg38 genome assembly using the Bowtie 2 software (40) version 2.3.4.1 with the parameters "--no-unal --no-discordant --no-mixed -sensitive-local". The mapped reads were filtered using the Samtools View software (41) version 1.13 with the parameters "-q 20 -b -F 0x04" to remove low-quality reads and duplicates. The resulting bam files were processed using the Deeptools bamCompare software version 3.5.4 (42), to create bigwig coverage files in ratio mode and normalized to RPKM with a minMappingQuality of 20 (6).

### Graphical profiling of protein occupancy across G4(+)

The occupancy of PQS, G4P, and proteins was quantitated across G4(+) with its bed file using the Deeptools computeMatrix software with the interval center as a reference point. Occupancy profile was obtained with the plotHeatmap command as described (6). The resulting PNG images were automatically cropped and contrast-normalized to 8-bit gray images using custom scripts by calling commands of the Imagemagick software (https://imagemagick.org/). The output images were then compared to a representative reference image using the -metric PAE option to assess their visual difference. Finally, the grayscale images were converted to pseudocolored images using a custom script by calling Imagemagick commands and a custom lookup table.

### Proteins

EGR1 was purchased from MedChemExpress LLC (Shanghai, China) and PCBP1, NONO from Detai Biotechnology (Nanjing, China).

### Electrophoretic mobility shift assay (EMSA)

EMSA was conducted as previously described (6) with modifications. DNAs (Table S1) were dissolved at 10 nM in a buffer containing 20 mM Tris-HCl (pH 7.4), 150 mM KCl, 1 mM EDTA,0.4 mg/ml BSA, denatured at 95 °C for 5 min, and slowly cooled down to 25 °C. Renatured DNA was then incubated with 250 nM cold DNA and 250 nM indicated protein at 4 °C for 20 min, followed by an addition of 10 nM FAM-labeled DNA and subsequent incubation at 4 °C for 40 min. DNA samples were resolved on 6 % non-denaturing polyacrylamide gel containing 50 mM KCl at 4 °C for 20 min in 1× TBE buffer. DNA was visualized on a ChemiDoc MP imager (Bio-Rad).

## Acknowledgments

This work was supported by the National Natural Science Foundation of China (grant # 22037004 and 22377009) and the Shanxi “1331 Project”.

## Supporting Information

**Table S1.**
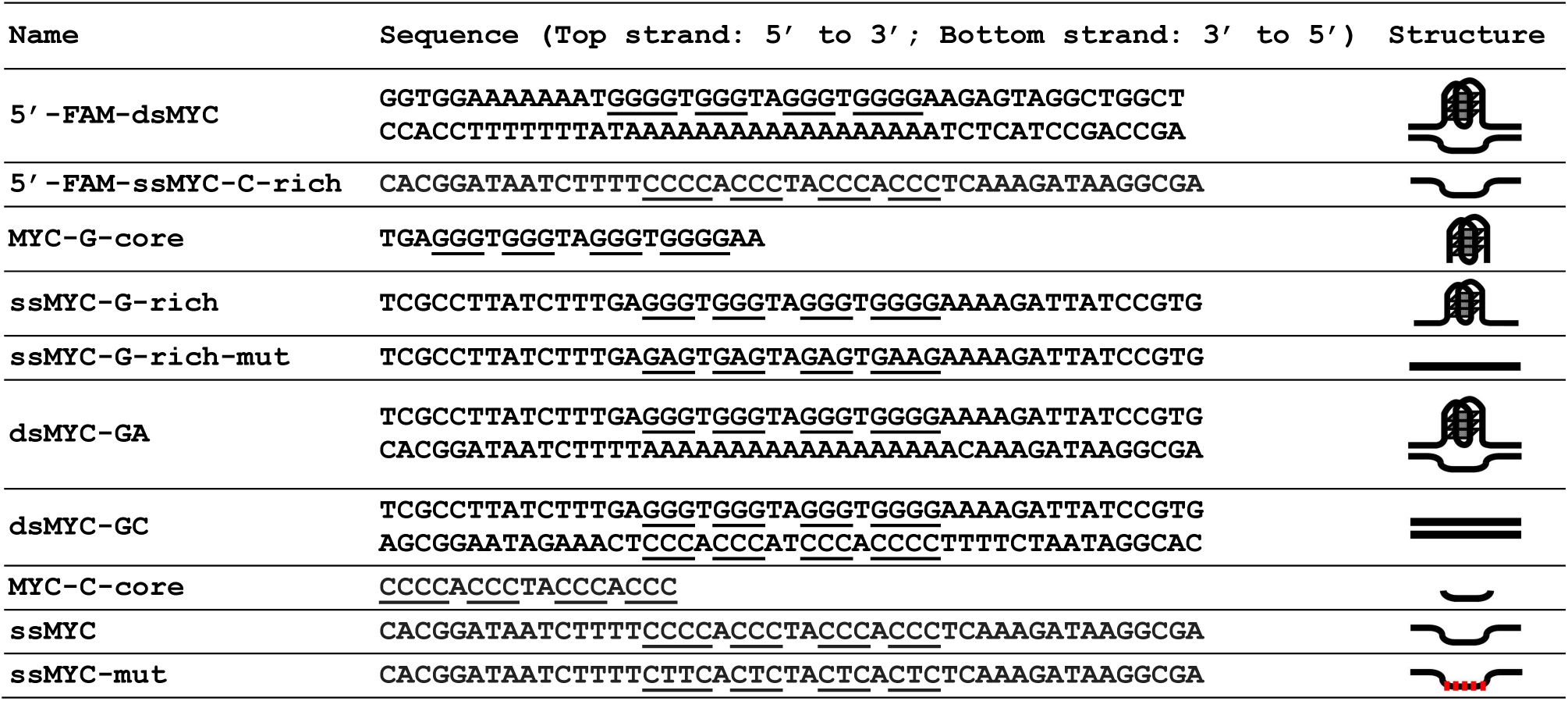
Sequences of DNAs used in EMSA.

**Figure S1.**
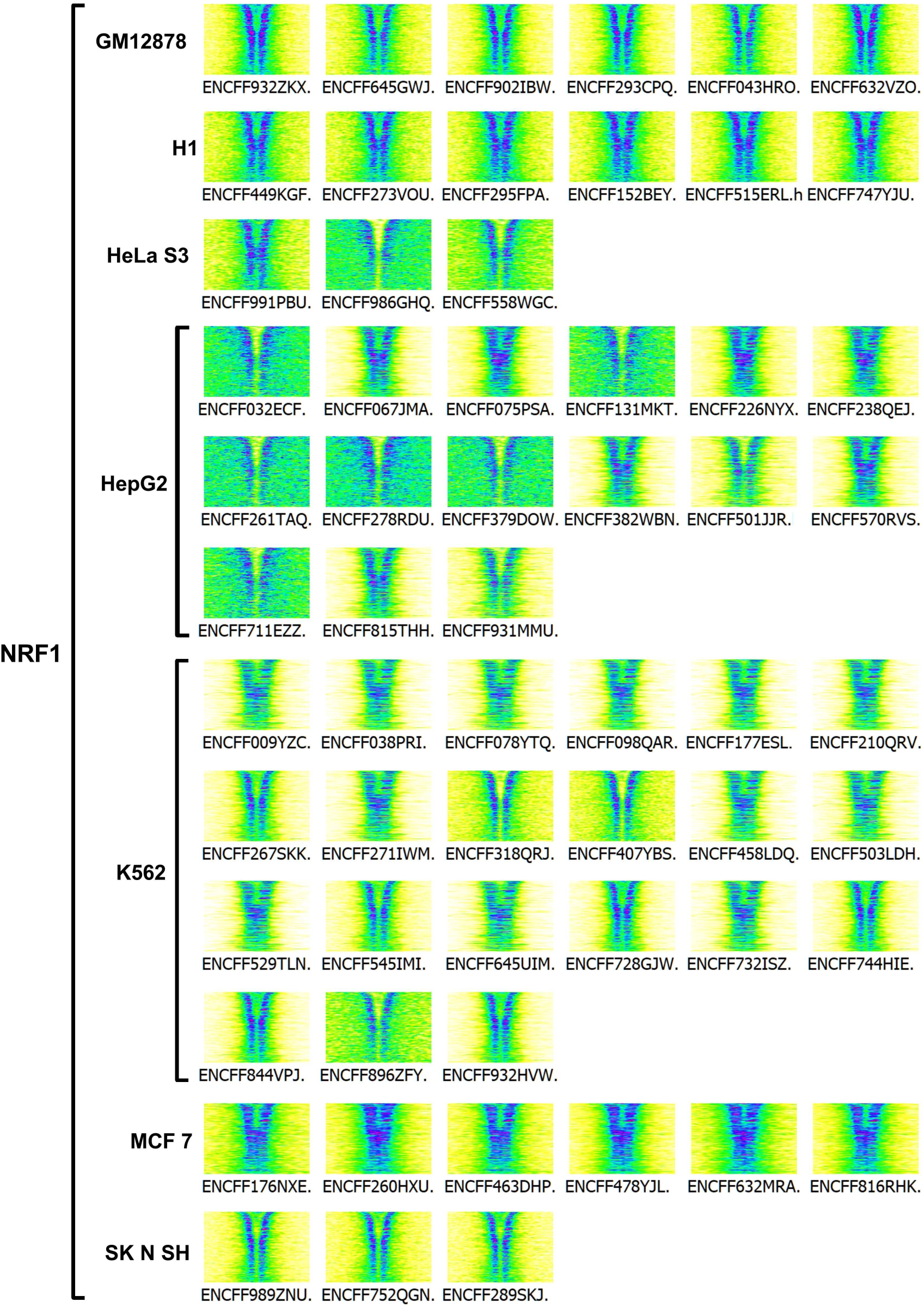
Occupancy of NRF1 around G4(+) in all samples from 7 cell lines showing consistent binding to G4(+) proximal side within and across cell types. Sample ID is shown below each image.

**Figure S2.**
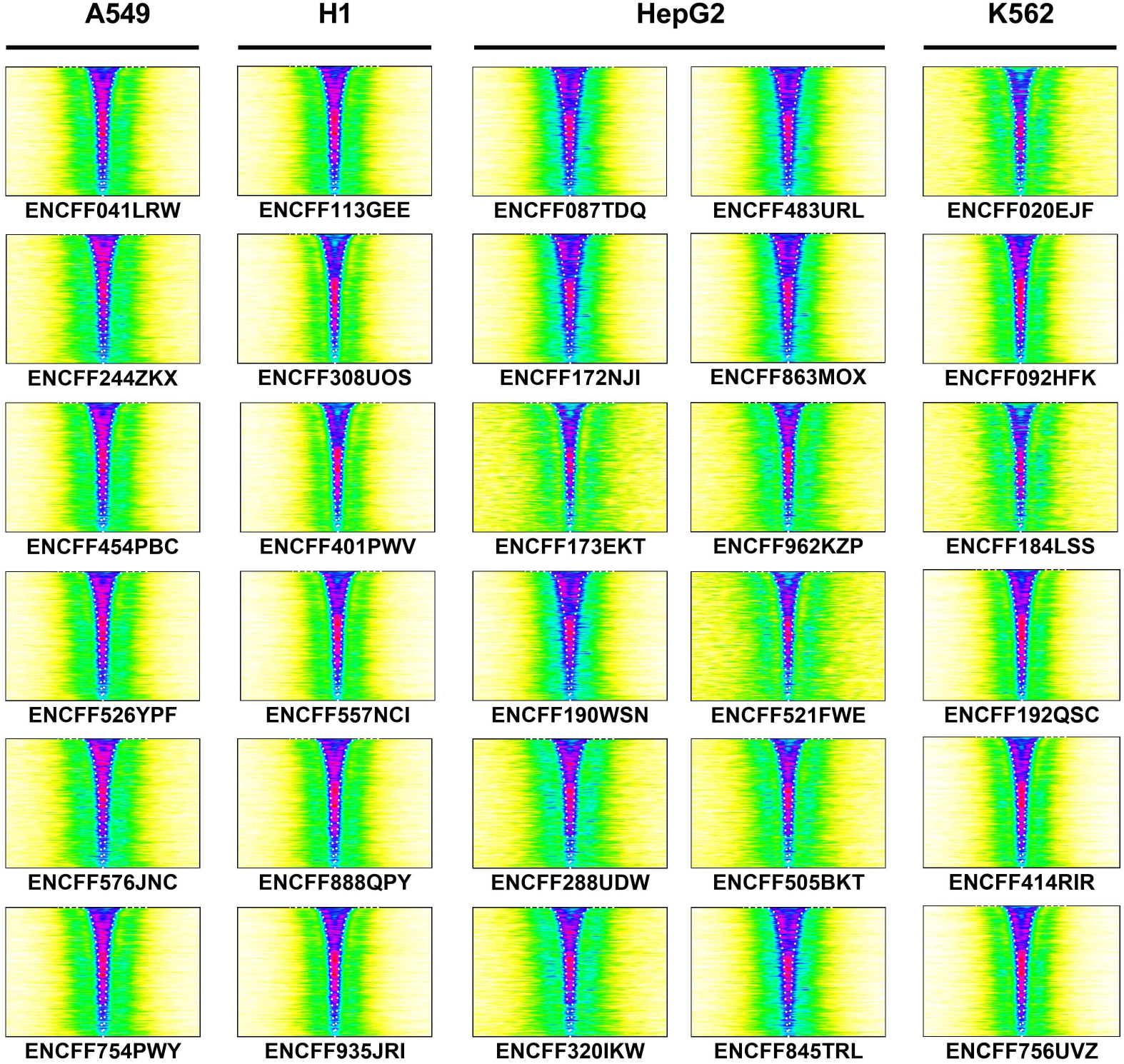
Occupancy of PHF8 in all samples from cell lines A549, H1, HepG2, and K562 showing consistent binding to G4(+) within and across cell types. Sample ID is shown below each image.

**Figure S3.**
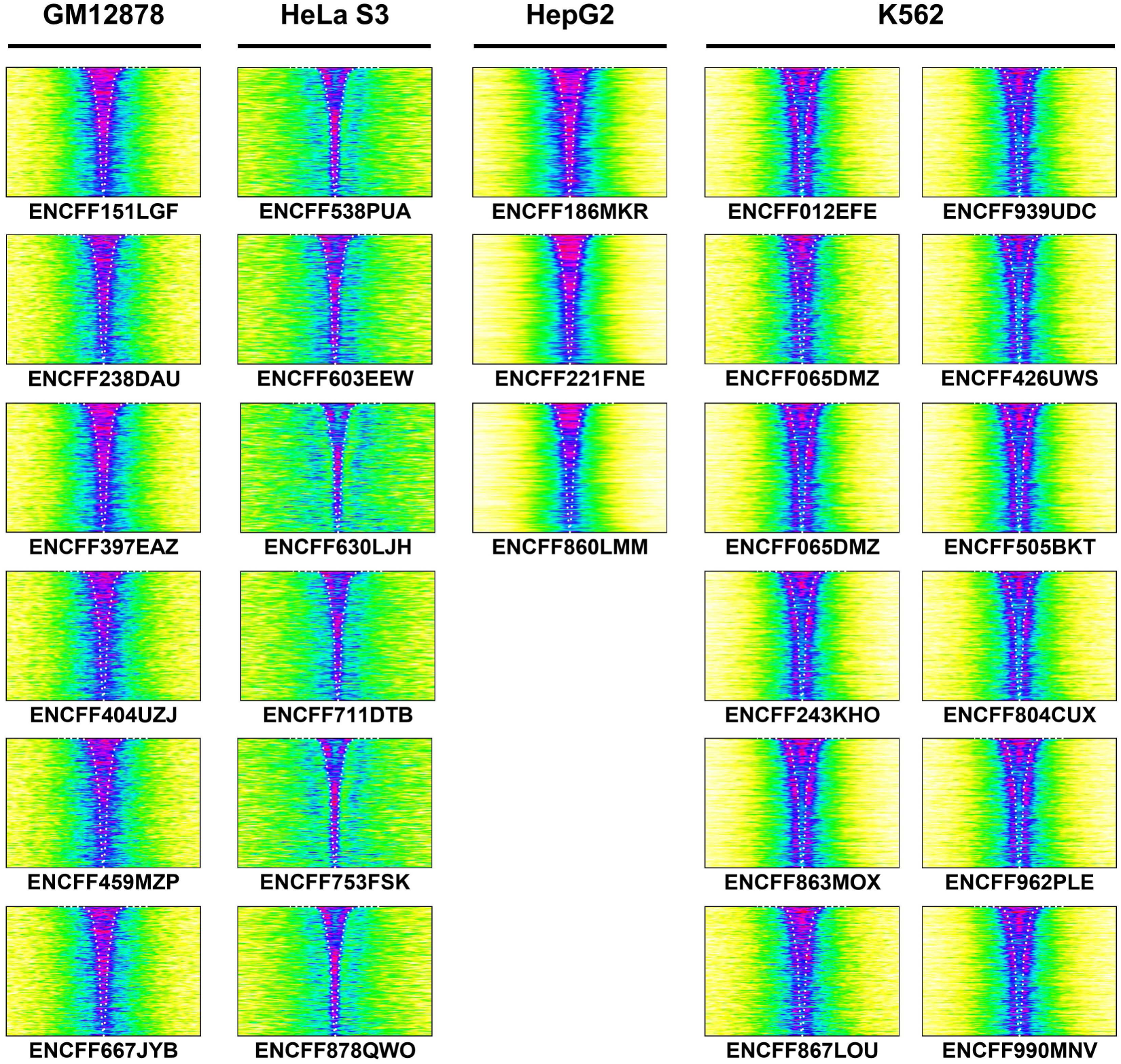
Occupancy of HBTF in all samples from cell lines GM12878, HeLa S3, HepG2, and K562 showing consistency within and variation between cell types. Sample ID is shown below each image.

**Figure S4.**
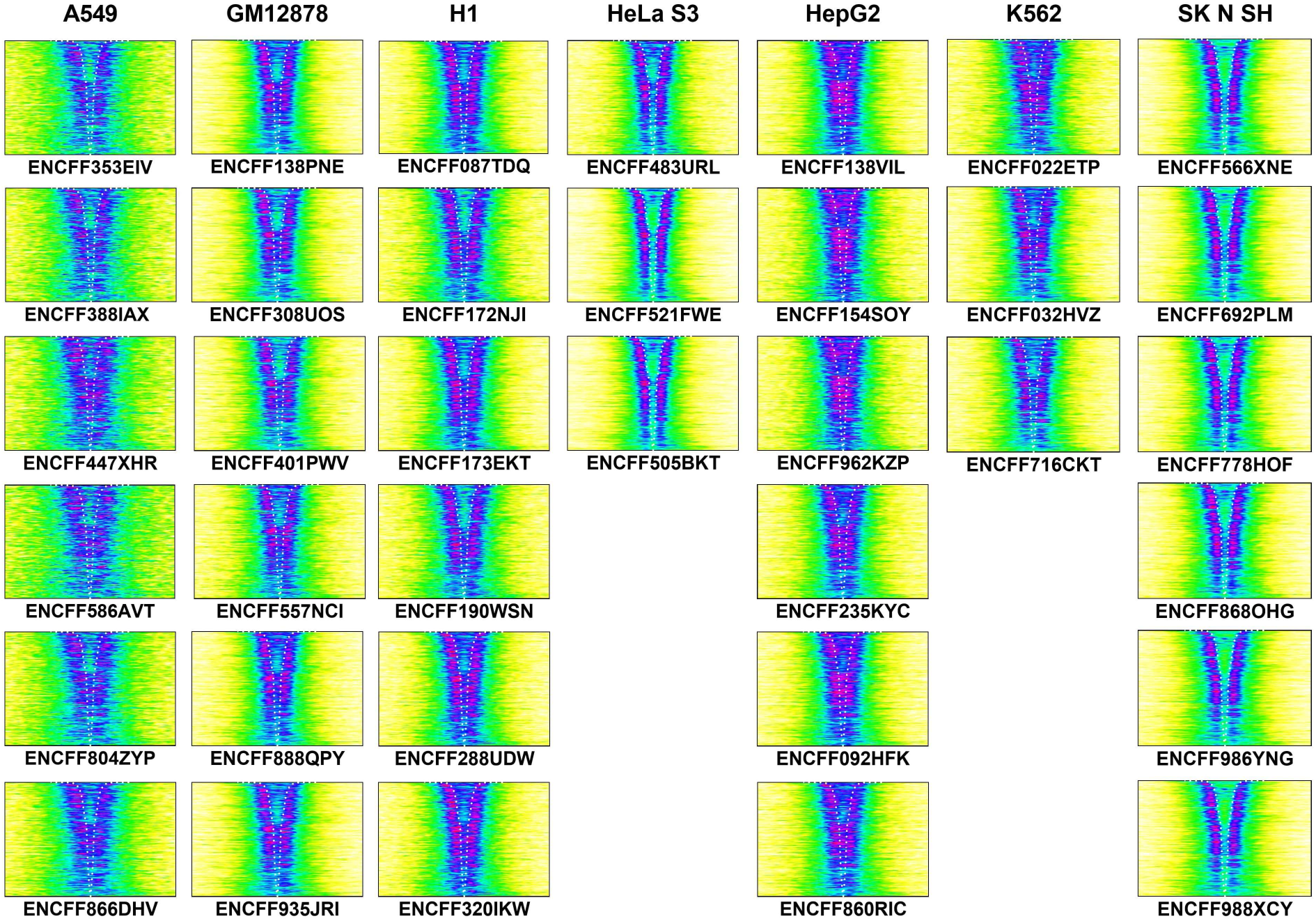
Occupancy of CHD2 in all samples from cell lines A549, GM12878, H1, HeLa S3, HepG2, K562, and SK N SH showing consistent binding to G4(+) proximal side within and across cell types. Sample ID is shown below each image.

**Figure S5.**
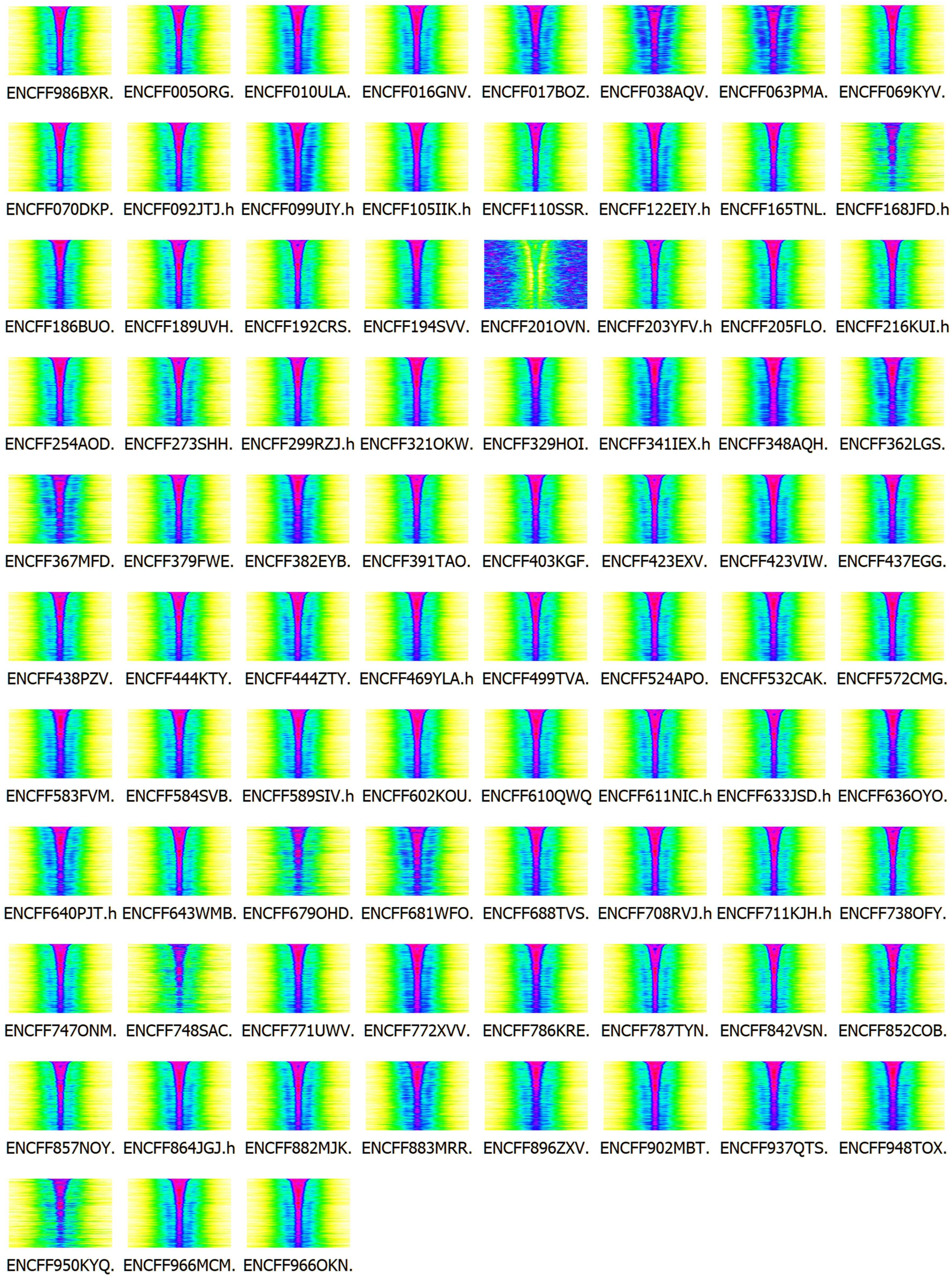
Occupancy of H3K27ac in all the 83 samples from cell line A549 showing consistent binding to G4(+) within the cell type. Sample ID is shown below each image.

**Figure S6.**
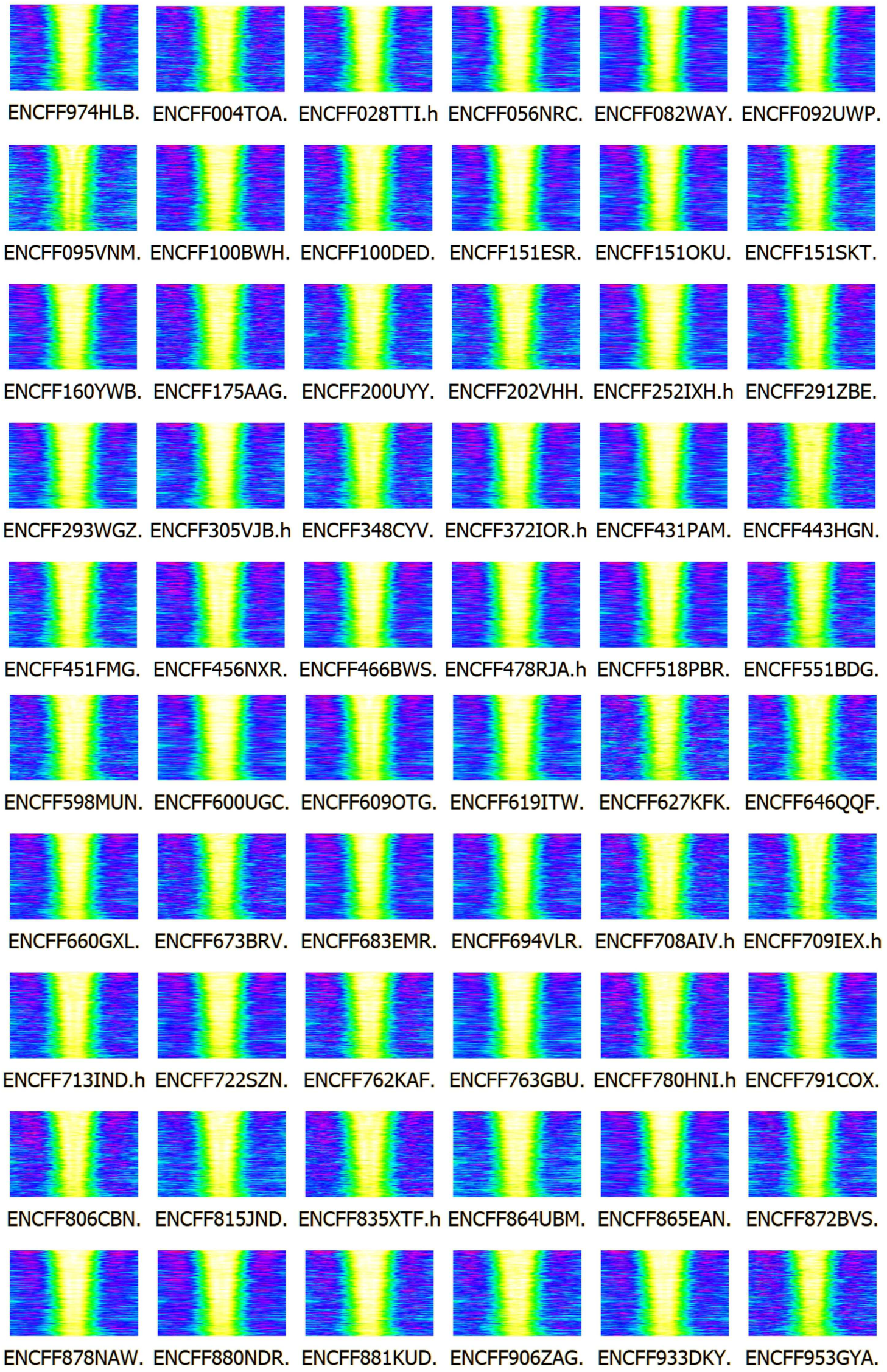
Occupancy of H3K4me1 in all the 60 samples from cell line A549 showing consistent depletion at G4(+) within the cell type. Sample ID is shown below each image.

**Figure S7.**
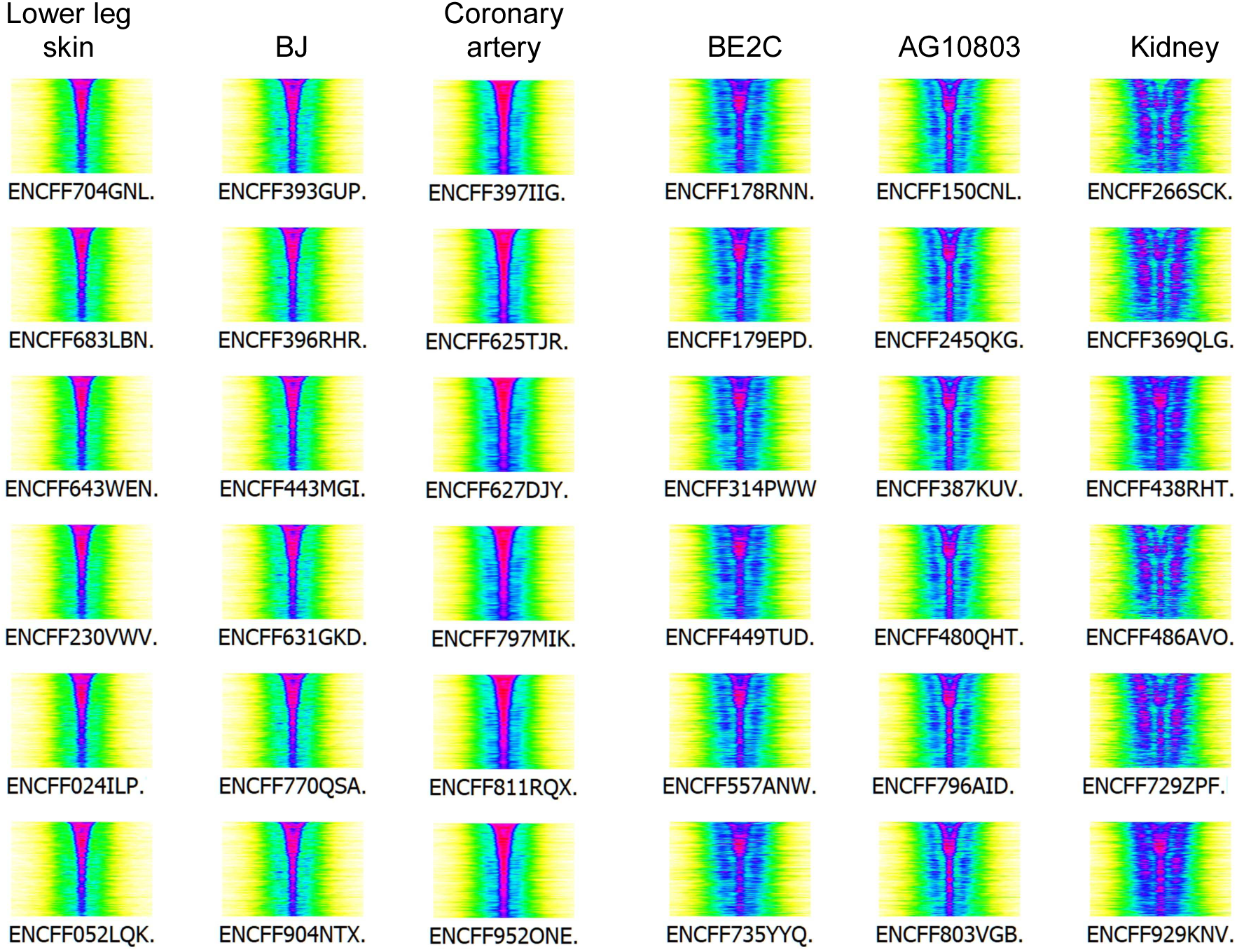
Occupancy of H3k4me3 in all samples from the indicated cell types, showing consistency within and variation between cell types. Sample ID is shown below each image.

**Figure S8.**
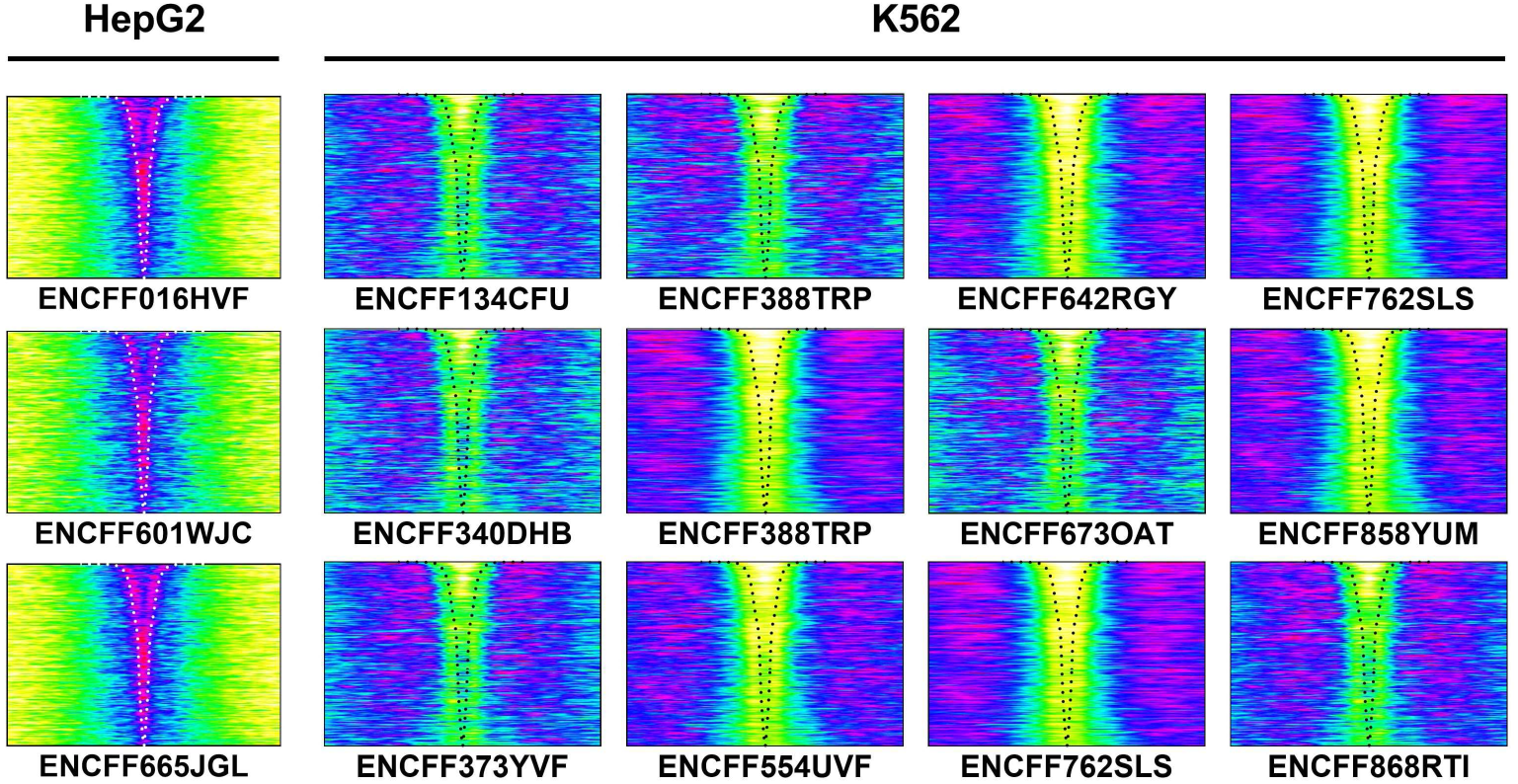
Occupancy of ZNF318 in all samples from cell lines HepG2 and K562 showing consistency within and variation between cell types. Sample ID is shown below each image.

**Figure S9.**
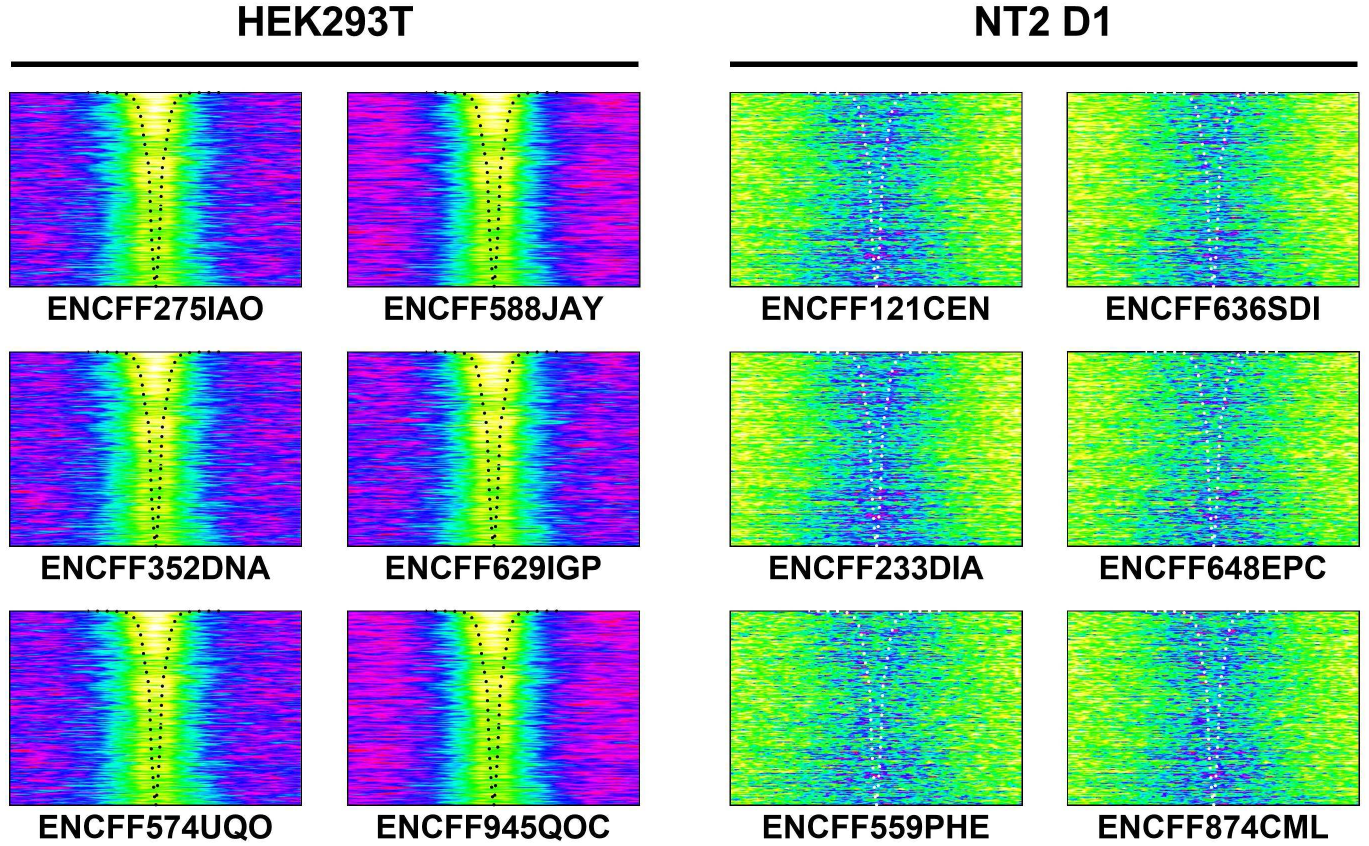
Occupancy of SUZ12 in all samples from cell lines HEK293T and NT2 D1 showing consistency within and variation between cell types. Sample ID is shown below each image.

**Figure S10.**
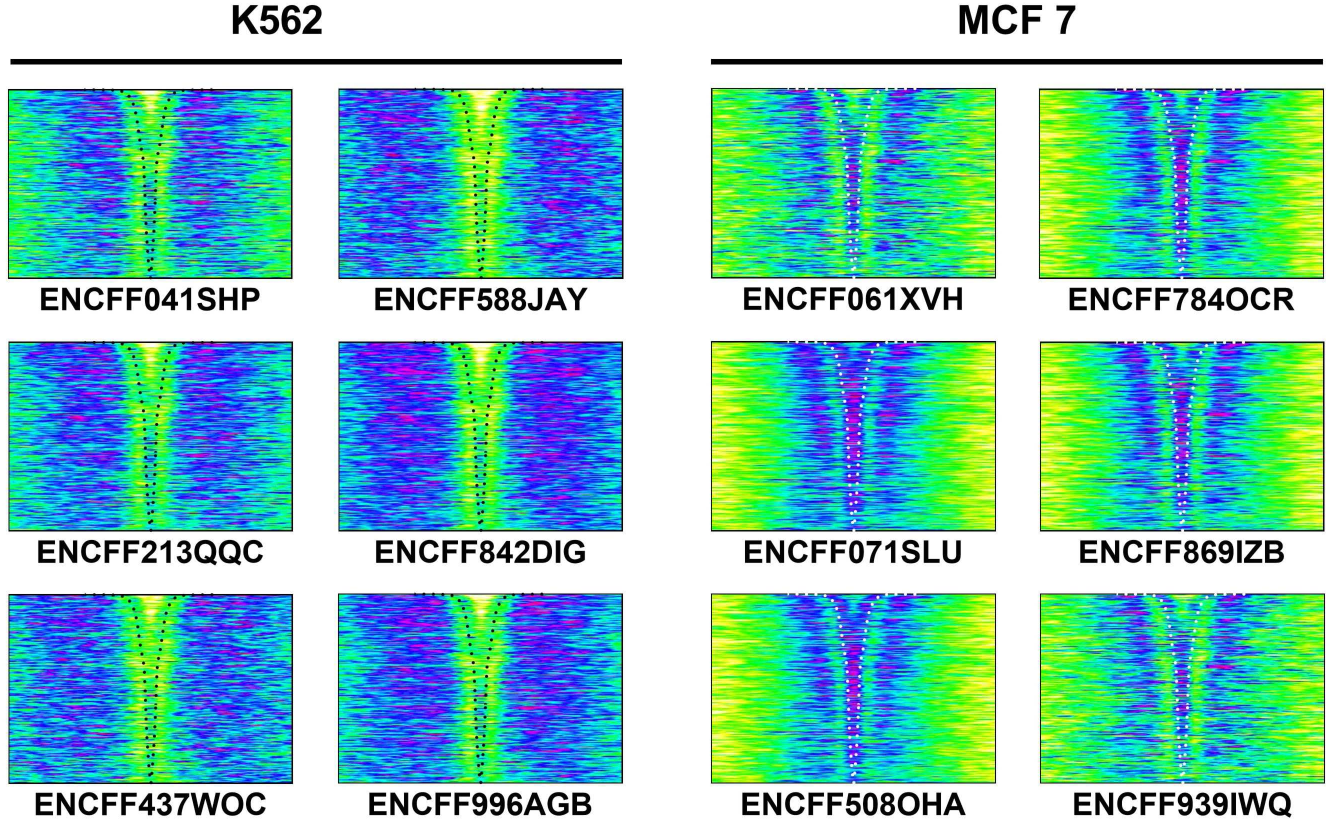
Occupancy of SNIP1 in all samples from cell lines K562 and MCF showing consistency within and variation between cell types. Sample ID is shown below each image.

**Figure S11.**
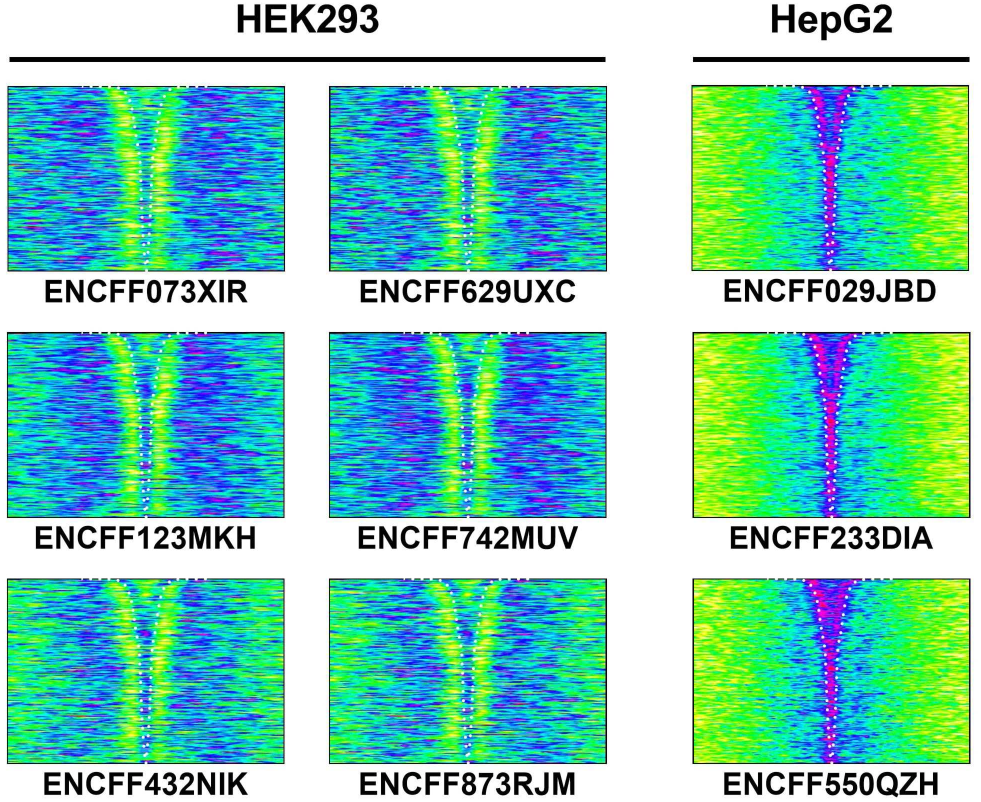
Occupancy of ZNF530 in all samples from cell lines HEK293 and HepG2 showing consistency within and variation between cell types. Sample ID is shown below each image.

**Figure S12.**
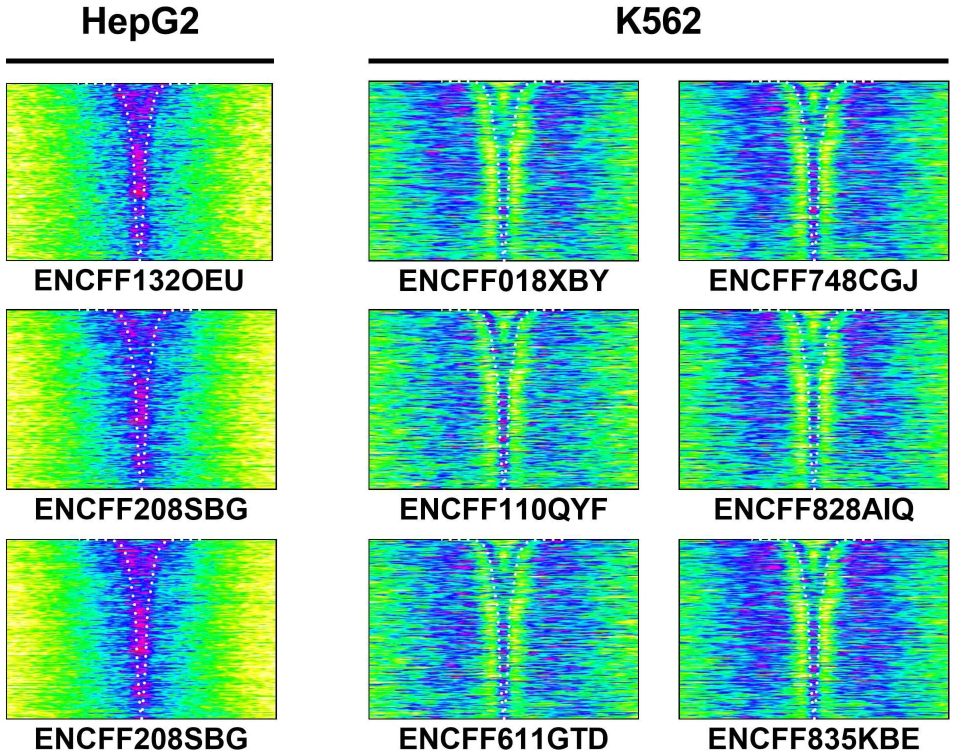
Occupancy of ARID2 in all samples from cell lines HepG2 and K562 showing consistency within and variation between cell types. Sample ID is shown below each image.

**Figure S13.**
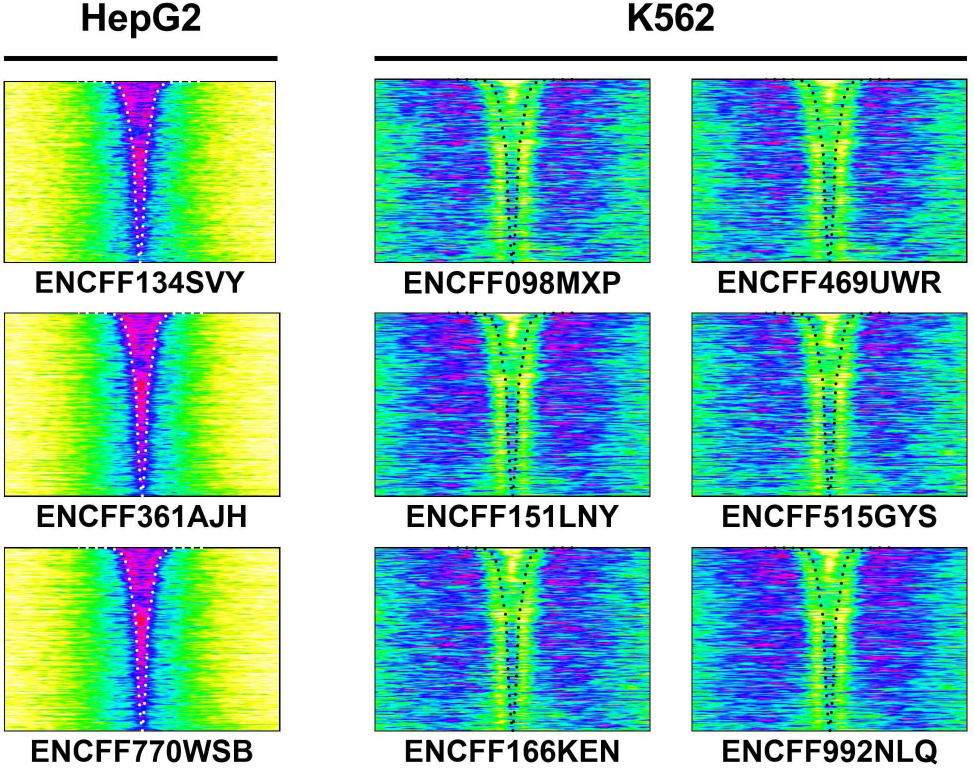
Occupancy of DNMT1 in all samples from cell lines HepG2 and K562 showing consistency within and variation between cell types. Sample ID is shown below each image.

**Figure S14.**
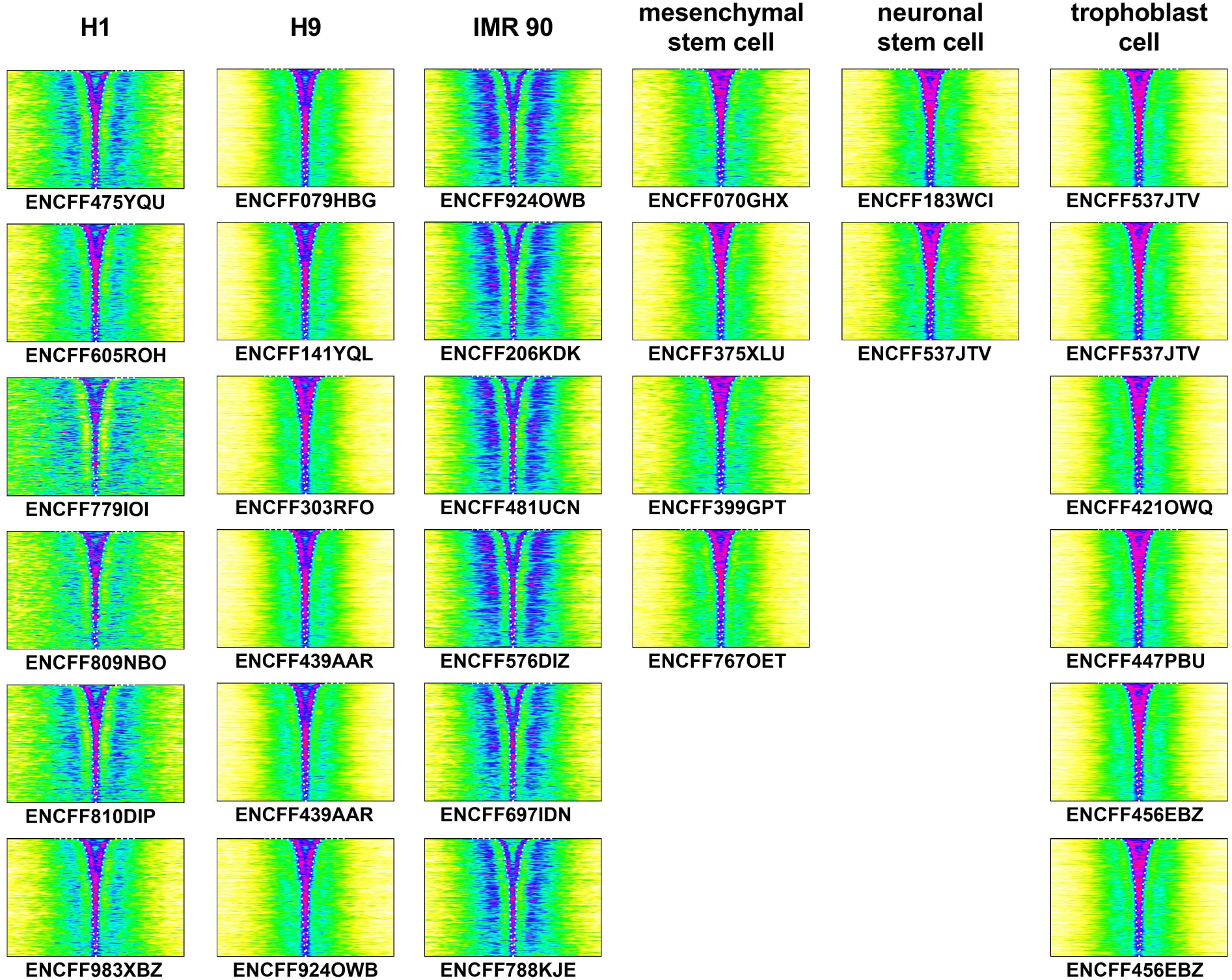
Occupancy of H3K14ac in all samples from cell lines H1, H9. IMR 90, in vitro differentiated mesenchymal stem cells, neuronal stem cells, and trophoblast cells showing consistency within and variation between cell types. Sample ID is shown below each image.

**Figure S15.**
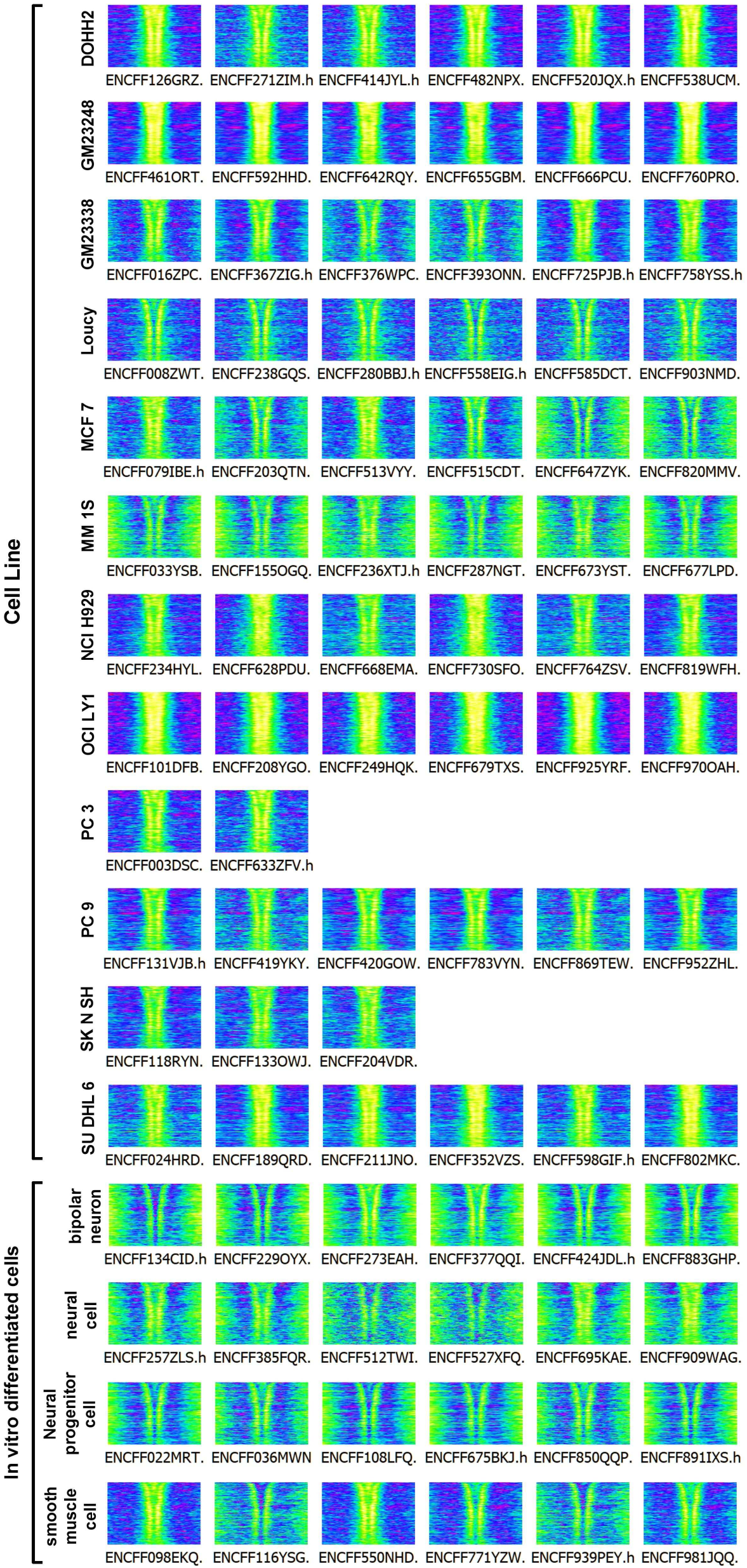
Occupancy of H3F3A in all samples from the indicated cell types showing consistency within and variation between cell types. Sample ID is shown below each image.

**Figure S16.**
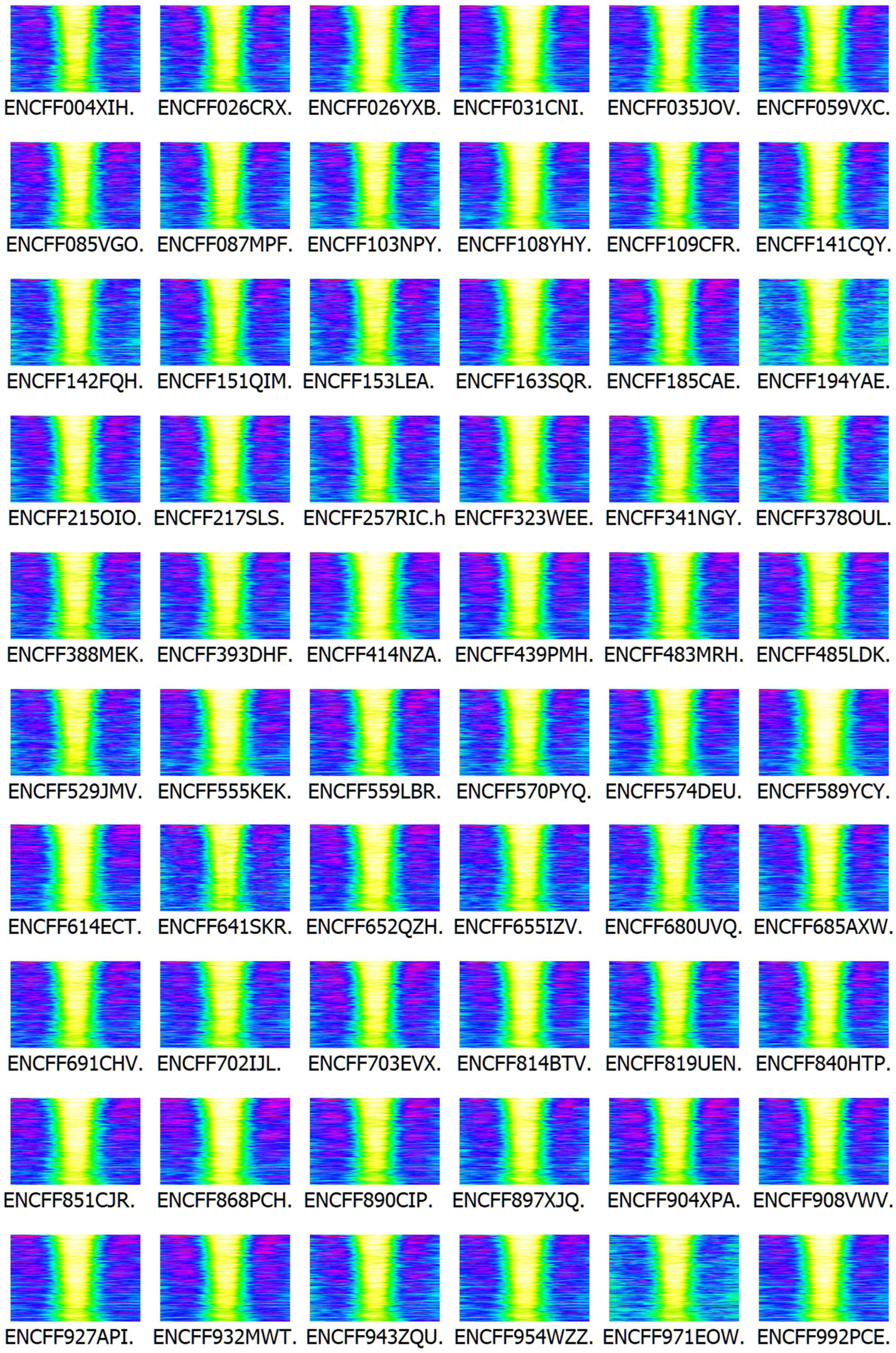
Occupancy of H3K4me1 in all the 60 samples from BLaER1 cells showing consistent depletion in G4(+) within cell type. Sample ID is shown below each image.

**Figure S17.**
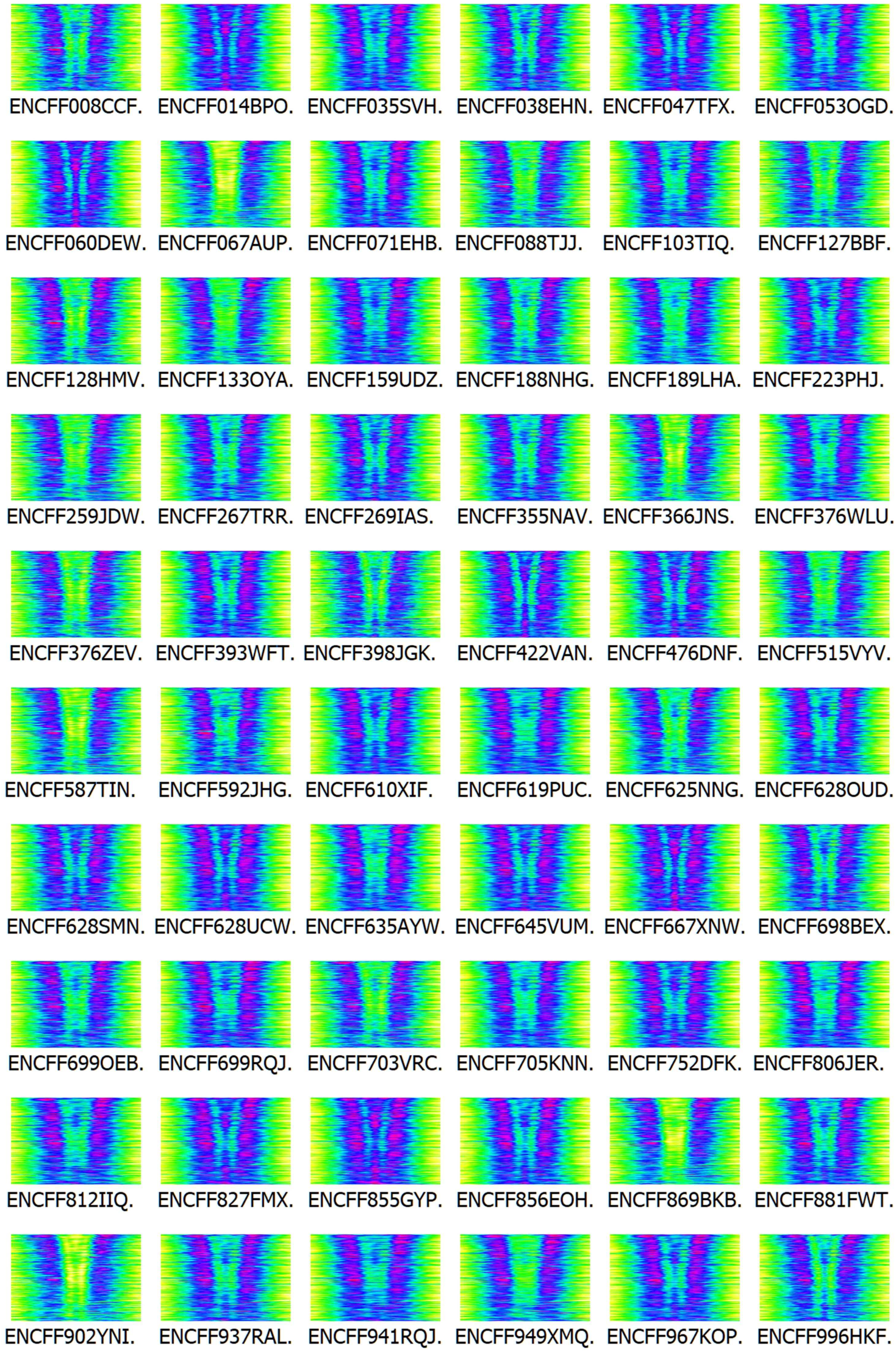
Occupancy of H3K4me2 in all the 60 samples from BLaER1 cells showing consistent binding at G4(+) distal side within the cell type. Sample ID is shown below each image.

**Figure S18.**
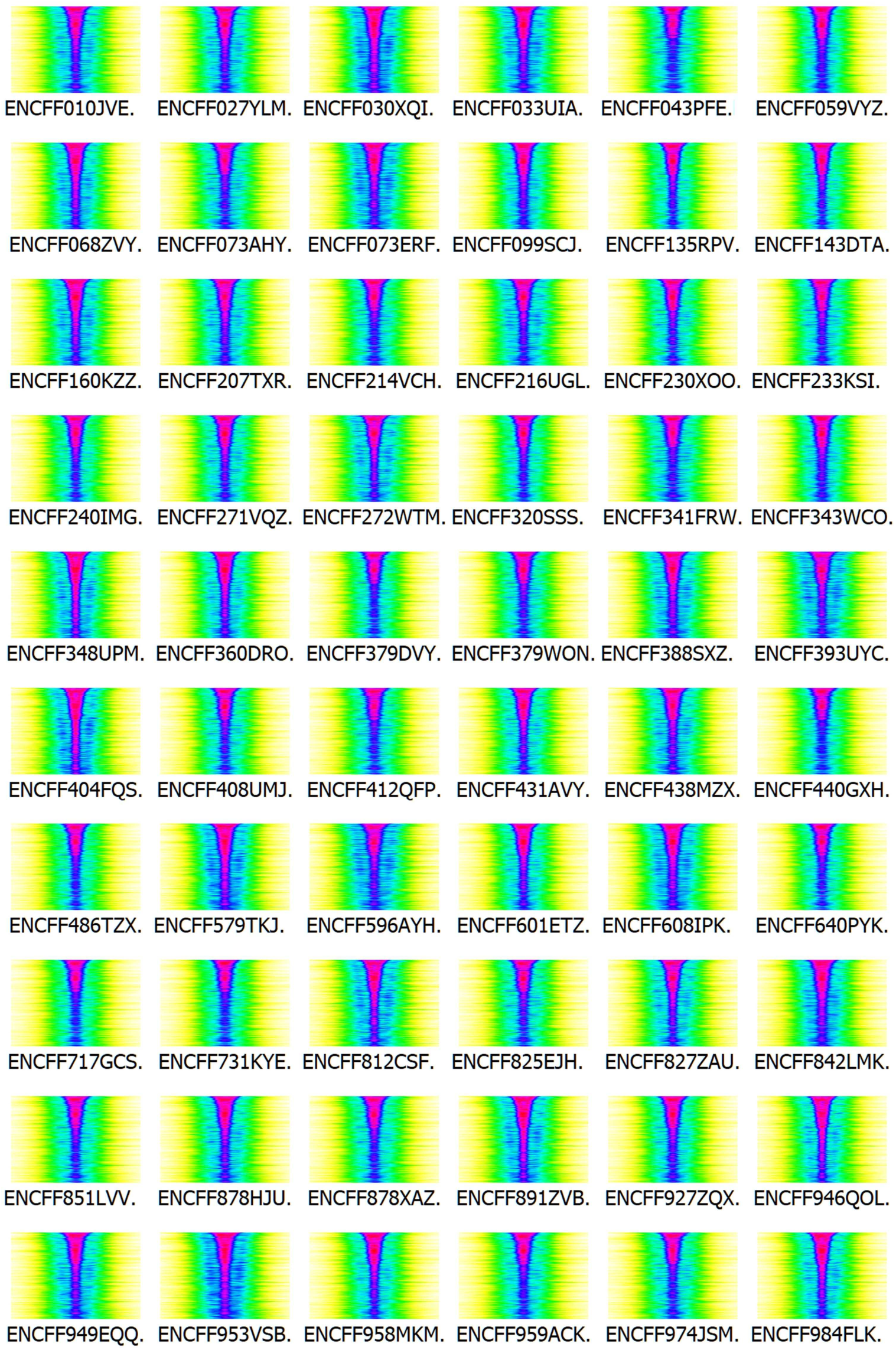
Distribution of H3K4me3 in all the 60 samples from BLaER1 cells showing consistent binding in G4(+) within the cell type. Sample ID is shown below each image.

**Figure S19.**
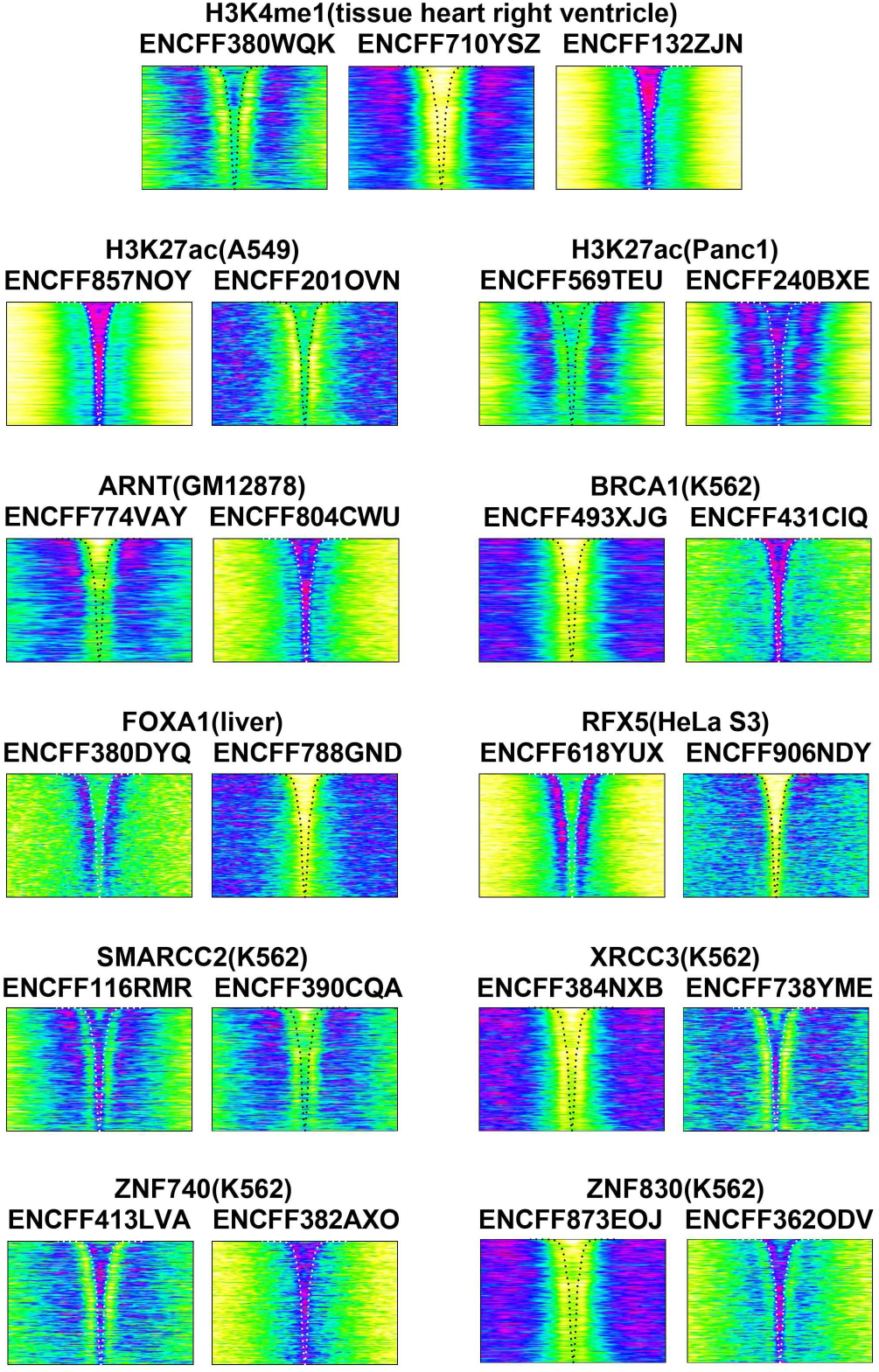
Examples of intra-cell type variation in protein occupancy across G4(+). Protein name, cell line, and ID are shown above each image.

